# Emotional states as distinct configurations of functional brain networks

**DOI:** 10.1101/2021.07.23.453552

**Authors:** Rotem Dan, Marta Weinstock, Gadi Goelman

## Abstract

The conceptualization of emotional states as patterns of interactions between large-scale brain networks has recently gained support. Yet, few studies have directly examined the brain’s network structure during emotional experiences. Here, we investigated the brain’s functional network organization during experiences of sadness, amusement, and neutral states elicited by movies, in addition to a resting-state. We tested the effects of the experienced emotion on individual variability in the brain’s functional connectome. Next, for each state, we defined a community structure of the brain and quantified its segregation and integration. We found that sadness, relative to amusement, was associated with higher modular integration and increased connectivity of cognitive control networks: the salience and fronto-parietal networks. Moreover, in both the functional connectome and the emotional report, the similarity between individuals was dependent on the sex. Our results suggest that the experience of emotion is linked to a reconfiguration of whole-brain distributed, not emotion-specific, functional networks and that the brain’s topological structure carries information about the subjective emotional experience.

## Introduction

The encoding of emotion in the human brain remains an area of ongoing debate. Traditionally, most studies sought to map specific categories of emotion to localized brain regions or anatomical circuits, focusing on the role of subcortical structures (Ekman 1999; Panksepp 2004). However, it has become increasingly evident that there is no one-to-one mapping between discrete emotions and individual brain regions (Lindquist et al. 2012). With the evolution of cognitive network neuroscience, the study of the neural basis of emotion has shifted to the examination of functional brain networks (Pessoa 2017). The recent application of machine learning techniques to neuroimaging data demonstrated that specific categories of emotion can be distinguished based on neural activity patterns, distributed over the cortex and subcortex (Kragel and LaBar 2015; Wager et al. 2015).

This new line of research may suggest that there is no brain region or even a brain network, that is specific to a certain emotion. Rather, emotional states are thought to be reflected in patterns of interactions between multiple large-scale brain networks (Barrett and Satpute 2013; Wager et al. 2015; Pessoa 2017). These functional brain networks are not only involved in emotion but also in non-emotional processes (such as conceptualization, attention, motor function, and executive control), and are not dedicated to an emotion category, but instead, their specific interactions are postulated to be the basis for an emotional state. However, surprisingly, despite much theoretical interest, few studies have directly examined the brain’s network structure during subjective experiences of emotion. Thus, in an attempt to fill these experimental gaps, our study’s goals were twofold. First, to delineate the manner in which emotional states influence individual differences in brain functional connectivity. Second, to define the community structure of the brain during specific emotional experiences.

Outside the field of emotion, the effects of the task or mental state on individual differences in the brain’s functional connectome have recently gained interest (Cole et al. 2013; Geerligs et al. 2015). A functional connectome refers to the communication patterns between brain regions, typically computed by the level of synchronization between fMRI blood-oxygen-level-dependent (BOLD) signals (Van Essen and Ugurbil 2012). The effects of the mental state on variability in the functional connectome can be evaluated by examining: (i) within-subject similarity, i.e., how functional connectomes acquired during different states are similar within an individual, and (ii) between-subject similarity, how functional connectomes acquired during a specific state are similar across individuals. Previous studies, such as that of Finn and colleagues (Finn et al. 2017), demonstrated that both the within- and between-subject similarities are dependent on the state.

The mental state was further shown to influence the brain’s modular organization. As a complex system, the brain can be divided into communities of regions, called modules, that are functionally highly intra-connected and less strongly functionally inter-connected (Meunier et al. 2010). Describing the brain from a network science perspective allows one to quantify higher-level complex interactions between the network’s elements (e.g., brain regions), and examine their link to the mental state and behavior (Medaglia et al. 2015). For example, Cohen and D’Esposito (Cohen and D’Esposito 2016) found increased modular segregation during a motor task as opposed to enhanced global integration during a working memory task.

Crucially, disturbances of emotion are key components in many psychiatric disorders, such as depression, anxiety, and autism. For developing brain-based models that can predict disease trajectory and clinical outcome, it is critical to understand how emotional states are encoded in the brain’s complex functional structure. Importantly, this includes not only the macro-scale brain networks that are involved in these states, but also the specific communication patterns between brain networks, and how they are dependent on the subjective emotional experience. Ultimately, such knowledge should advance the clinical utility of neuroimaging methods.

In the current study, we aimed to: (i) Examine the effects of the emotional experience and sex on the within- and between-subject similarity in the brain’s functional connectome. We hypothesized that the emotional state would increase the similarity in the brain’s global organization, both between individuals and between states within individuals. Given the well-established sex differences in mood disorders (Riecher-Rössler 2017), we hypothesized that sex would interact with the emotional state. (ii) Define the modular organization of the brain during specific emotions: sadness and amusement. These emotional categories were chosen based on their relevance to clinical disorders, such as depression and bipolar disorder (Joormann et al. 2007; Fu et al. 2008; Gruber et al. 2014), and their opposed emotional valence. (iii) Quantify differences between emotional states in the segregation and integration of network communities. In line with the idea of emotional states as distinct interactions between large-scale functional brain networks, we postulated that sadness and amusement, despite being both high-order complex cognitive states, will differ in their level of modular segregation and in specific patterns of between-module communication.

## Materials and Methods

### Participants

Healthy young participants were recruited among undergraduate students at the Hebrew University of Jerusalem. Exclusion criteria included: past or present psychiatric or neurological disorders, use of psychiatric medication, use of hormonal contraceptives, or any premenstrual symptoms. One participant was excluded due to anxiety in the MRI scanner, and another was excluded because of in-scanner motion (see MRI data preprocessing in the Supplemental Methods). The final analyzed sample comprised 50 healthy participants (30 women, 20 men; age: 23.94±2.64 years). Datasets from a subsample of the participants were previously used in a prior study (Dan et al. 2019). For women that completed two MRI scans as part of a separate parallel study, data utilized in this study were always taken from their first scan. Note that half of the women were scanned during the mid-follicular phase and half during the late-luteal phase. The study was approved by the Hadassah Hebrew University Medical Center Ethics Committee and was carried out in compliance with the Declaration of Helsinki.

### Emotional brain states induced by movie-watching

Each fMRI scan included the following sequence to induce four different mental states: (i) resting-state (10 minutes): eyes open, fixating on a visual crosshair; (ii, iii) two emotional states (10 minutes each): sadness and amusement, induced by continuous exposures to movies, and (iv) a neutral movie state (10 minutes). Each movie state (sadness, amusement, and neutral) was induced by four movie clips (2-3 minutes each) presented in a row. The order of the emotional states (sadness, amusement) was counterbalanced across the sample and within each sex. The resting-state preceded the movie states and the neutral state was induced between the emotional states. We chose movies since we wanted to probe emotion in a naturalistic way, with an intense emotional experience. Movie-watching is a powerful ecological method to induce engaging and rich high-order cognitive states in the MRI scanner (Nastase et al. 2020). Another advantage of movie-watching is that it commonly reduces in-scanner head motion and thus can improve data quality (Vanderwal et al. 2019).

The movies were taken from available sets (Gross and Levenson 1995; Rottenberg et al. 2007; Farb et al. 2010; Schaefer et al. 2010) and tested in a separate behavioral study on an independent group of participants to evaluate their discreetness, intensity, and to select the best movie clips to use in the scanner. The following criteria were applied (Dan et al. 2019): (i) the movies should induce only the target discrete emotion (e.g., sadness and not a mixture of sadness and fear) (ii) the target emotion should be rated high on intensity. The movies were matched for the duration, the number of actors, and social interaction. All movies were in English with Hebrew subtitles. Detailed information about the movie stimuli is provided in Supplemental Table S1.

Before the MRI scan, participants were instructed to allow themselves to feel whatever emotions that come up without trying to suppress them. Within the scanner, participants rated after each movie state their subjective emotional experience during movie-watching on 1-to-8 Likert scales for: sadness, amusement, fear, anger, calmness, arousal, and attention. Responses were collected using MRI-compatible button boxes with 4 for each hand (8 buttons total). Outside the scanner, participants rated their subjective emotional experience during each movie state on 1-to-8 Likert scales using a detailed questionnaire that included 17 discrete emotional categories, valence (i.e., the affective quality of the experience, good/positive vs. bad/negative), arousal, and interest. The questionnaire, translated to Hebrew (see Supplemental Methods for the English version) was similar to that of Gross and colleagues (Rottenberg et al. 2007) and participants were encouraged to report honestly their emotions and try to separate them from their mood that day, from what they think other people felt, or what they believe people should feel.

### MRI data acquisition and preprocessing

MRI data were acquired on a 3T Magnetom Skyra. Preprocessing of fMRI data was done using SPM12 (Wellcome Trust Centre for Neuroimaging, London, United Kingdom, http://www.fil.ion.ucl.ac.uk/spm/software/spm12). Details of acquisition, preprocessing, and in-scanner motion are found in the Supplemental Methods.

### Functional connectome for each brain state

For each participant and mental state (i.e., sadness, amusement, neutral, rest), a whole-brain functional connectome covering the cortex and subcortex (excluding the cerebellum) was computed using CONN (Whitfield-Gabrieli and Nieto-Castanon 2012). The brain was parcellated to nodes and the average fMRI BOLD time series was calculated for each node. Pearson’s correlations were computed between time series from all pairs of nodes and Fisher’s transform was applied to correlation values, resulting in symmetric connectivity matrices.

In a network analysis of the brain, the effect of the atlas used for node definition is an area of ongoing investigation. We, therefore, calculated functional connectomes using four different atlases: (i) Automated anatomical labeling atlas (AAL) (Tzourio-Mazoyer et al. 2002): 90 nodes; (ii) Harvard-Oxford atlas (HO) (Desikan et al. 2006): 105 nodes; (iii) Shen atlas (Shen et al. 2013): 218 nodes (excluding the cerebellum); (iv) Schaefer atlas (Schaefer et al. 2018): 400 nodes. The brain stem and cerebellum were excluded from the atlases. Since the Schaefer brain subdivision does not include the subcortex, 14 subcortical nodes from the Harvard-Oxford atlas were added to this atlas, resulting in a total of 414 nodes. Note that the AAL and Harvard-Oxford subdivisions were defined based on anatomical features whereas Shen and Schaefer defined parcels based on the functional homogeneity of resting-state time series.

### Functional connectome similarity analysis

For each parcellation atlas, functional connectivity matrices were thresholded at z=0 (i.e., negative weights were set to zero) to include only positive correlations. To minimize the potential confounding effects of visual features of the movie stimuli on the state-specific functional connectivity matrices, the entire occipital cortex was excluded from the functional connectome similarity analyses. While auditory features may also have confounding effects, the impact of the auditory cortex was considered less substantial relative to the occipital cortex, as the latter includes approximately 15% of parcels in a brain atlas. Of note, recent studies that reconstructed movie stimuli from whole-brain fMRI signals have shown that the decoders rely most heavily on the visual cortex, and especially on low-level visual areas (Wang et al. 2022).

To examine within- and between-subject similarity in the functional connectome, we followed previous approaches (Cole et al. 2014; Finn et al. 2017): the unique elements of each connectivity matrix were extracted by taking the upper triangle of the matrix, resulting in a vector of edge values for each participant for each mental state. The between-subject similarity was calculated using Pearson’s correlation between all possible pairs of participants for a certain state. These values were Fisher’s z-transformed, averaged across participants, and reverted to r values, resulting in a single value per participant per state. The within-subject similarity was calculated using Pearson’s correlation between all pairs of states for each participant, resulting in six values per participant (sadness vs. amusement, sadness vs. neutral, amusement vs. neutral, sadness vs. rest, amusement vs. rest, neutral vs. rest).

To examine the effect of sex on between-subject similarity, we calculated similarity among women (i.e., between each woman and all other women) and among men, denoted here as women-to-women and men-to-men similarity. To verify that the unbalanced number of women and men in the sample (30 women, 20 men) did not create a bias, we repeated the sex differences analyses with 20 men and a randomly chosen subsample of 20 women. Statistical analyses were conducted with SPSS v.23 (SPSS Inc., Chicago, IL) on the Fisher transformed z-values. Differences in within- and between-subject similarity were tested by 2-way mixed ANOVA models implemented by a GLM with state as a within-subject factor and sex as a between-subject factor. Post-hoc pairwise comparisons were conducted using Sidak correction.

### Modularity analysis

#### Community detection for each brain state

To decompose the entire brain network into functional communities for each mental state, modularity analysis was conducted on the weighted functional connectivity matrices using a consensus partitioning algorithm (Lancichinetti and Fortunato 2012) implemented in the Brain Connectivity Toolbox (BCT) (Rubinov and Sporns 2010). This algorithm partitions a whole-brain network into nonoverlapping groups of brain regions, i.e., “communities” or “modules”, by maximizing a modularity function. The Harvard-Oxford atlas was chosen for node definition since it showed greater between-subject similarity in the functional connectome per state, compared to the higher resolution network parcellations of Shen and Schaefer (see Results and Figure 2a). In the first step, for each participant and each state separately, we applied a Louvain modularity optimization procedure 500 times to define an agreement matrix. Each cell in the agreement matrix corresponded to the proportion of times that a pair of nodes were assigned to the same community over the 500 repetitions. At the next step, Louvain modularity optimization was conducted on the agreement matrix, with 500 repetitions, generating a new agreement matrix. This step was repeated until convergence, i.e., until the agreement matrix was binary: containing only zeros or ones.

**Figure 1.**
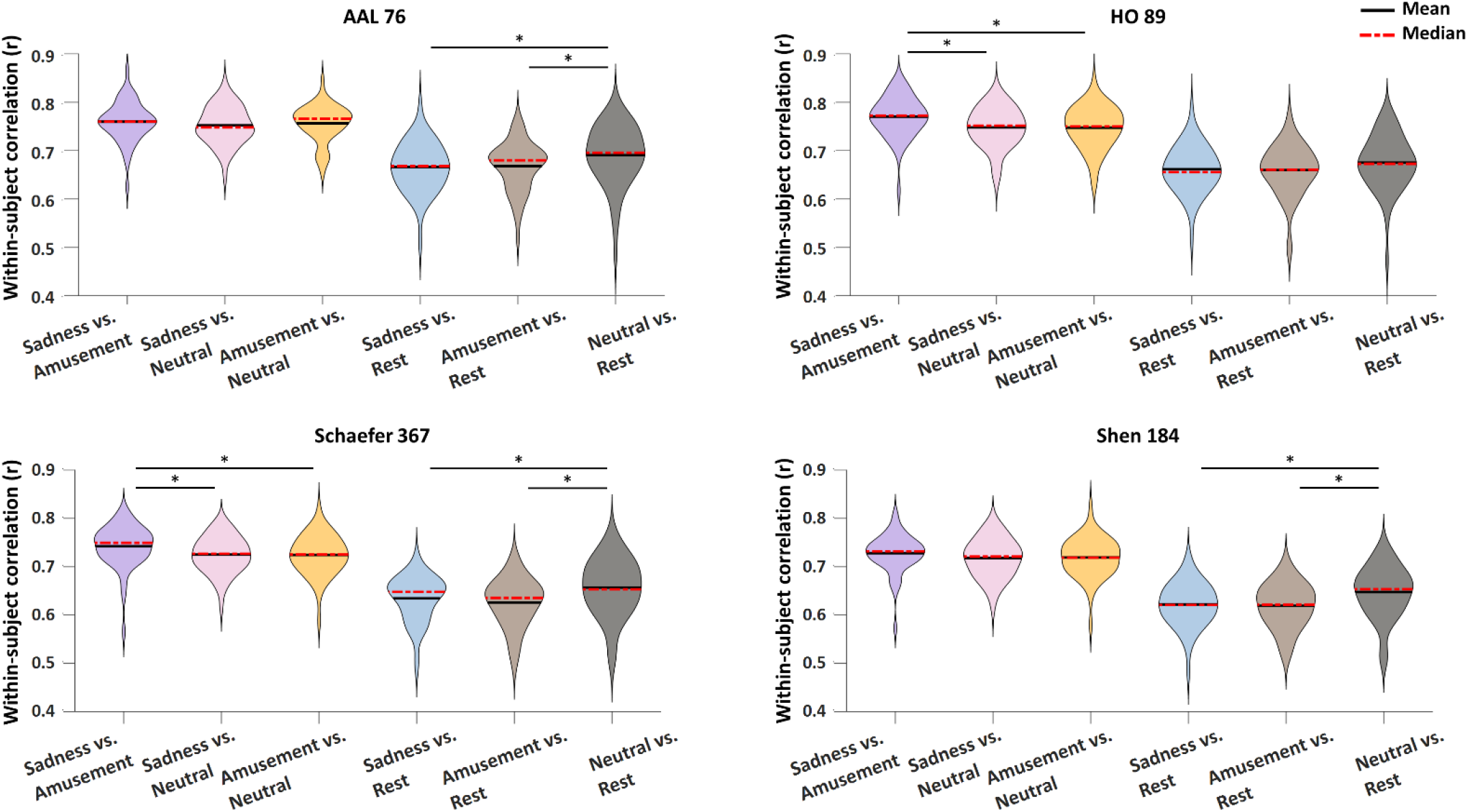
Within-subject similarity in the functional connectome is dependent on the brain state. The within-subject similarity in the functional connectome (Pearson’s correlation r values) is presented as a function of the pair of brain states, for the four parcellation atlases. The number of regions is indicated next to the atlas’s name. In each violin, the median is indicated by dashed red lines and the mean by solid black lines. Significant differences are marked by asterisks (*), *p*<0.05 Sidak corrected. Note that all pairwise differences between the 3 most left violins and the 3 most right violins were significant, and not marked by lines and asterisks for visual clarity.

**Figure 2.**
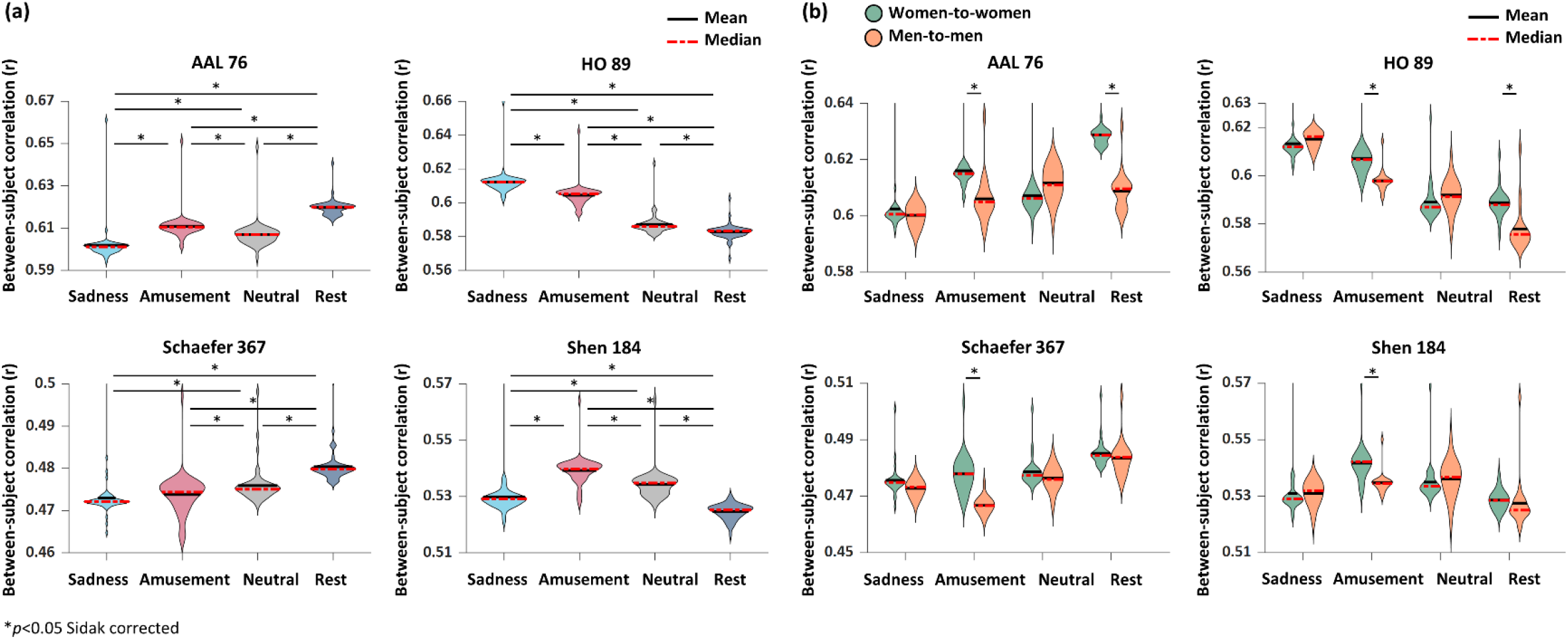
Between-subject similarity in the functional connectome is dependent on the brain state, sex, and parcellation atlas. The between-subject similarity in the functional connectome (Pearson’s correlation r values) is presented as a function of the brain state, for the four parcellation atlases. The number of regions is indicated next to the atlas’s name. **(a)** The between-subject similarity was dependent on the state and the brain parcellation atlas. **(b)** The effect of sex was examined by computing women-to-women (green) and men-to-men (orange) similarity. A significant sex-by-state interaction was found for all parcellation atlases. In each violin, the median is indicated by dashed red lines and the mean by solid black lines. Significant differences are marked by asterisks (*), *p*<0.05 Sidak corrected.

After a consensus partition was identified for each participant, we computed a group-level consensus partitioning for each state. The consensus partitions of all participants for a specific state were combined to create a group-level agreement matrix. Each cell in the group-level agreement matrix corresponded to the proportion of times that a pair of nodes were assigned to the same community across participants. At the next step, Louvain modularity optimization was conducted on the group-level agreement matrix, using 500 repetitions, generating a new group-level agreement matrix. This step was repeated until convergence to a binary group-level matrix. The procedure yielded a community structure, or modular organization, for each state at the individual and group levels. The consensus partitioning procedure was repeated over a range of the resolution parameter gamma (Reichardt and Bornholdt 2006) and with different treatments of negative weights in the modularity function (Rubinov and Sporns 2011).

A resolution parameter gamma=2.25 and a positive-only weighted modularity function were chosen. Details regarding the choice of the resolution parameter and modularity function are found in the Supplemental Methods, Figures S1-S5.

#### Modular segregation and integration metrics

After each node was assigned to a group-level module for each mental state, we calculated the following measures of modular segregation and integration. Note that all measures retain the weights of the functional connections and were calculated for a specific state. Similarly to the functional connectome analyses, visual modules were excluded to minimize the possible confounding impact of the visual movie stimuli.

i. System segregation (Chan et al. 2014): the normalized difference between within- and between-module connectivity. A higher value indicates greater segregation, i.e., less interaction between modules. This measure was calculated for each module, resulting in a modular segregation metric as follows:

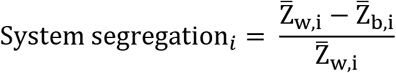

where 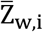 is the mean functional connectivity (Fisher transformed z-values) between all pairs of nodes within module i and 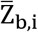 is the mean functional connectivity between nodes of module i and nodes of all other modules.
ii. Participation coefficient (Guimerà and Amaral 2005): quantifies the diversity of a node’s connections across modules. It is close to 1 if the node’s connections are distributed uniformly among all modules and 0 if all connections are within the node’s module. A high participation coefficient indicates high integration, i.e., strong connections to many modules, whereas a low participation coefficient indicates high segregation, i.e., weak interactions with other modules. For each module, we calculated the mean participation coefficient across all its nodes. The weighted variant of the participation coefficient from the BCT was used:

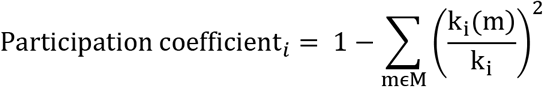

where *M* is the set of modules *m, k_i_*(*m*) is the weighted degree (i.e., summed weighted functional connectivity) of node i to nodes within module *m* and *k_i_* is the total weighted connectivity of node i to all nodes regardless of module membership.
iii. Pairwise between-module connectivity: defined as the mean functional connectivity strength (Fisher transformed z-values) between pairs of modules, averaged across all possible connections. For example, the connectivity between module i and module j was calculated as the sum of functional connections between nodes in module i and nodes in module j, divided by the number of possible connections (number of nodes in module i x number of nodes in module j).

To examine differences between the four mental states in overall modular segregation and integration, the mean system segregation (across all modules, excluding visual) and the mean participation coefficient (across all nodes, excluding visual) were computed and analyzed by 1-way ANOVA models. Statistical differences between the emotional states (sadness, amusement) in the segregation and integration of specific modules, were tested by 2-way repeated-measures ANOVA models implemented by a GLM with emotion and module as within-subject factors. Post-hoc pairwise comparisons were conducted using Sidak correction. In addition, for each emotional state, the association between segregation and integration metrics and the reported valence, arousal, and intensity of the emotional experience was computed using Pearson’s correlation. Ratings were taken from outside the scanner.

#### Behavioral report of emotion experience

Valence and interest were rated outside the scanner for the three movie states: sadness, amusement, and neutral. Arousal and the intensity of the target emotion were rated both within and outside the scanner. For the detailed questionnaires outside the scanner, the intensity of sadness was calculated as the average rating for “sadness” and “sorrow” and the intensity of amusement was calculated as the average of “amusement” and “enjoyment”. Attention was rated within the scanner. Note that arousal was added to the questionnaires (within and outside the scanner) after the beginning of the study; thus ratings were missing from 9 participants for all states. The rating of valence for amusement was missing from one participant.

The between-subject similarity in the behavioral report of emotion was computed for each movie state (sadness, amusement, neutral), utilizing a similar approach to the one used for the functional connectome. The detailed ratings outside the scanner were used and the vector of ratings for the 19 items (emotional categories, valence, interest) was extracted for each participant for each state. The between-subject similarity was calculated using Pearson’s correlation between all possible pairs of participants for a certain state. These values were Fisher’s z-transformed and averaged across participants resulting in a single value per participant. To examine the effect of sex, we calculated similarity among women and men, i.e., women-to-women and men-to-men similarity. To verify that the unbalanced number of women and men in the sample (30 women, 20 men) did not create a bias, we repeated the sex differences analysis with 20 men and a randomly chosen subsample of 20 women.

For each behavioral measure (valence, arousal, intensity, interest, attention, between-subject similarity), statistical analysis was conducted in SPSS by 2-way mixed ANOVA models implemented by a GLM with state as a within-subject factor and sex as a between-subject factor. Post-hoc pairwise comparisons were conducted using Sidak correction.

#### Discreteness of the target emotional state

Discreetness of the target emotional state was analyzed for both the ratings within and outside the scanner. For the detailed questionnaires completed outside the scanner, a score was computed for each of the following 11 emotion categories: sadness (calculated as the average rating for “sadness” and “sorrow”), amusement (average of “amusement” and “enjoyment”), fear, anger, embarrassment (average of “embarrassment” and “shame”), disgust, surprise, guilt, anxiety, contempt, and confusion. To evaluate whether each movie state induced a discrete target emotion, Wilcoxon signed-rank paired tests were computed for each state between the intensities of the target emotion (sadness or amusement) and all other 10 non-target emotional categories. Similarly, for the within-scanner ratings, Wilcoxon signed-rank paired tests were computed for each state between the intensities of the target emotion and the other 3 non-target emotional categories (fear, anger, sadness/amusement). A Bonferroni correction was applied for multiple comparisons (2 movie states, 13 comparisons for each movie) resulting in a significance threshold of *p*<1.9·10^-3^.

#### Quantifying visual and auditory features of movie stimuli

Low- and mid-level visual and auditory features of the movie stimuli were extracted using *Pliers* open-source Python package (McNamara et al. 2017) (https://github.com/PsychoinformaticsLab/pliers). The low-level features of brightness, vibrance, optic flow, and audio root-mean-square energy (audio RMSE) were extracted, as well as the mid-level feature of the number of faces presented onscreen. Note that for greater accuracy, the detection of faces was done both in *Pliers* and by manual encoding. The following metrics were computed: (i) *Pliers:* the proportion of frames containing at least one face, (ii) manual encoding: the number of faces presented for each frame. To assess potential differences between movie states in audiovisual features, for each movie clip and each feature, the mean and standard deviation across frames were calculated. For each feature, differences between movie states (sadness, amusement, neutral) in the mean across frames or in the standard deviations were assessed in SPSS by 1-way ANOVA models.

## Results

### Visual and auditory features of movie stimuli

Supplemental Table S2 presents the visual and auditory features of each movie clip. There were no statistical differences between states (sadness, amusement, neutral) for any of the movie features.

### Within-subject similarity in the brain’s functional connectome

The within-subject similarity was dependent on the state (Figure 1) [main effect of state, all *p*<10^-6^, Greenhouse-Geisser corrected: AAL: *F*(3.43,165.07)=81.84, partial eta-squared: η_p_^2^=0.63; HO: *F*(3.72,178.75)=106.62, η_p_^2^=0.69; Shen: *F*(3.70,178.02)=139.12, η_p_^2^=0.74; Schaefer: *F*(3.53,169.84)=158.30, η_p_^2^=0.76].

An individual’s functional connectome was more similar between two movie states (sadness, amusement, neutral) than between a movie state and rest, regardless of the brain parcellation atlas (pairwise differences: all *p*<10^-6^ Sidak corrected, Cohen’s *d*=0.99-2.92) (for visual clarity, this effect is not marked by lines and asterisks in Figure 1). For all atlases except for the HO, the functional connectome was more similar between resting-state and the neutral movie state than between resting-state and an emotional state (sadness or amusement) (all *p*<0.03 Sidak corrected, Cohen’s *d*=0.47-0.87). Furthermore, for the HO and Schaefer atlases, the functional connectome was more similar between the two emotional states (sadness, amusement) than between an emotional state and the neutral movie state (all *p*<0.03 Sidak corrected, Cohen’s *d*=0.46-0.56). There was no effect of sex or sex-by-state interaction (Supplemental Figure S6). The same results were found for a subsample matched for the number of women and men (20 women, 20 men), namely no sex differences (Supplemental Figure S7).

### Between-subject similarity in the brain’s functional connectome

Parcellating the brain into more nodes resulted in an overall decrease in the similarity between participants (number of parcels: Schaefer>Shen>HO>AAL) (Figure 2). The between-subject similarity was dependent on the state (Figure 2a) [main effect of state, all *p*<10^-6^ Sidak corrected, Greenhouse-Geisser corrected: AAL: *F*(1.79,88.01)=276.42, η_p_^2^=0.84; HO: *F*(2.16,105.86)=844.32, η_p_^2^=0.94; Shen: *F*(1.63,80.16)=201.22, η_p_^2^=0.80; Schaefer: *F*(1.70,83.47)=90.28, η_p_^2^=0.64]. In addition, the between-subject similarity was dependent on the parcellation atlas: for the HO and Shen atlases, participants were more similar to one another during movie-watching compared to rest, whereas an opposite effect was found for the AAL and Schaefer atlases, where participants were more similar to one another during rest compared to movie-watching.

Sex differences varied by state (Figure 2b) [sex-by-state interaction: AAL:*F*(1.87,89.84)=62.61, *p*<10^-6^, η_p_^2^=0.56; HO: *F*(1.81,87.02)=29.72,*p*<10^-6^, η_p_^2^=0.38; Shen: *F*(1.62,77.90)=9.02, *p*=10 ^-3^, η_p_^2^=0.15; Schaefer: *F*(2.51,120.49)=24.06, *p*<10^-6^, η_p_^2^=0.33; all Greenhouse-Geisser corrected]. Amusement showed significant sex differences across all atlases, with greater similarity among women (all *p*<10^-3^ Sidak corrected, Cohen’s *d*=0.55-0.96). Similar results were obtained with a balanced subsample of 20 women and 20 men (Supplemental Figure S8).

### Behavioral report of emotional experience

Table 1 presents the ratings of the movie states for discrete emotion categories, valence, arousal, interest, and attention, rated within and outside the scanner.

**Table 1.** Ratings of the movie states for discrete emotion categories, valence, arousal, interest, and attention.

|  | Sadness movie state |  | Neutral movie state |  | Amusement movie state |  |
| --- | --- | --- | --- | --- | --- | --- |
|  | (mean±std) |  | (mean±std) |  | (mean±std) |  |
|  | Outside scanner | Within scanner | Outside scanner | Within scanner | Outside scanner | Within scanner |
| Sadness | **5.89±1.54** | **6.02±1.74** | 1.20±0.61 | 1.30±0.70 | 1.19±0.57 | 1.20±0.45 |
| Amusement | 2.66±1.44 | 2.94±1.88 | 2.68±1.30 | 3.18±1.57 | **6.15±1.34** | **6.40±1.14** |
| Fear | 2.22±1.76 | 2.98±2.10 | 1.02±0.14 | 1.16±0.37 | 1.16±0.65 | 1.56±1.12 |
| Anger | 2.00±1.44 | 2.30±1.48 | 1.20±0.63 | 1.36±0.82 | 1.24±0.51 | 1.38±0.69 |
| Disgust | 1.18±0.48 | - | 1.12±0.38 | - | 1.78±1.20 | - |
| Surprise | 1.50±1.12 | - | 1.12±0.38 | - | 3.16±1.95 | - |
| Embarrassment | 1.29±0.76 | - | 1.20±0.56 | - | 2.32±1.35 | - |
| Guilt | 1.26±0.75 | - | 1.00±0 | - | 1.14±0.63 | - |
| Anxiety | 1.80±1.34 | - | 1.00±0 | - | 1.28±0.63 | - |
| Contempt | 1.42±0.88 | - | 1.54±1.01 | - | 1.68±1.05 | - |
| Confusion | 1.68±1.33 | - | 1.30±0.78 | - | 1.36±0.63 | - |
| Calmness | 3.68±1.94 | 3.58±1.95 | **5.19±1.87** | **5.74±1.74** | 4.92±1.92 | 5.46±1.74 |
| Valence | 4.28±1.79 | - | 5.36±1.36 | - | 6.77±1.10 | - |
| Arousal | 5.63±1.75 | 5.41±1.84 | 3.29±1.74 | 3.53±1.91 | 6.04±1.39 | 6.09±1.68 |
| Interest | 5.02±1.93 | - | 2.88±1.57 | - | 5.78±1.65 | - |
| Attention | - | 6.46±1.22 | - | 3.78±1.60 | - | 6.46±1.38 |

Within the scanner, ratings were done on 1-to-8 Likert scales and collected immediately after each movie state. Participants made their responses using MRI-compatible button boxes with 4 for each hand (8 buttons total). Outside the scanner, ratings were done with pen and paper on 1-to-8 Likert scales. The intensity of the target emotion is indicated in bold.

### Valence, arousal, intensity, interest, and attention

#### (i) Valence

A main effect of state was found, as expected: amusement>neutral>sadness [*F*(2,94)=53.46, *p*<10^-6^, η_p_^2^=0.53]. Namely, the emotional experience during induction of amusement was rated as “pleasant/positive” whereas the experience during induction of sadness was rated as “unpleasant/negative”. There were no sex or interaction effects.

#### (ii) Arousal

A main effect of state was found [(i) within scanner: *F*(2,78)=40.66, *p*<10^-6^, η_p_^2^=0.51; (ii) outside scanner: *F*(1.63,63.75)=47.45, *p*<10^-6^, η_p_^2^=0.54, Greenhouse-Geisser corrected]. Higher arousal was reported for both emotional states relative to the neutral state. Outside the scanner, no difference was found between emotional states. Within the scanner, higher arousal was reported for amusement relative to sadness (*p*=0.041 Sidak corrected, Cohen’s *d*=0.36). There were no sex or interaction effects.

#### (iii) Intensity of target emotion

Within or outside the scanner, emotional states (sadness, amusement) did not differ in the reported intensity of target emotion, and there was no effect of sex or interaction.

#### (iv) Interest

Higher interest was reported for amusement relative to sadness, and for both emotional states relative to neutral: amusement>sadness>neutral [main effect of state: *F*(2,96)=47.10, *p*<10^-6^, η_p_^2^=0.49]. There were no sex or interaction effects.

#### (v) Attention

Higher attention was reported during emotional states relative to the neutral state, without a difference between emotional states [main effect of state: *F*(1.72,67.4)=64.28, *p*<10^-6^, η_p_^2^=0.62, Greenhouse-Geisser corrected]. There were no sex or interaction effects.

### Discreteness of the target emotional state

For both emotional states, all pairwise comparisons between the intensities of the target emotion and other non-target emotion categories were significant (all *p*<10^-7^), across ratings collected outside and within the scanner, indicating discrete emotional experiences of sadness and amusement (Supplemental Table S3).

### Between-subject similarity in the behavioral report

Participants were more similar to one another in their reports of emotional experience during amusement, relative to sadness or the neutral state (Figure 3a) [main effect of state: *F*(2,98)=2774.61, *p*<10^-6^, η_p_^2^=0.98]. In addition, women were more similar to other women than men to other men (Figure 3b) [main effect of sex: *F*(1,48)=17.20, *p*=1.36·10^-4^, η_p_^2^= 0.26. main effect of state: *F*(2,96)=1236.42, *p*<10^-6^, η_p_^2^=0.96]. There was no sex-by-emotion interaction. Similar results were obtained with a balanced subsample of 20 women and 20 men (Supplemental Figure S9).

**Figure 3.**
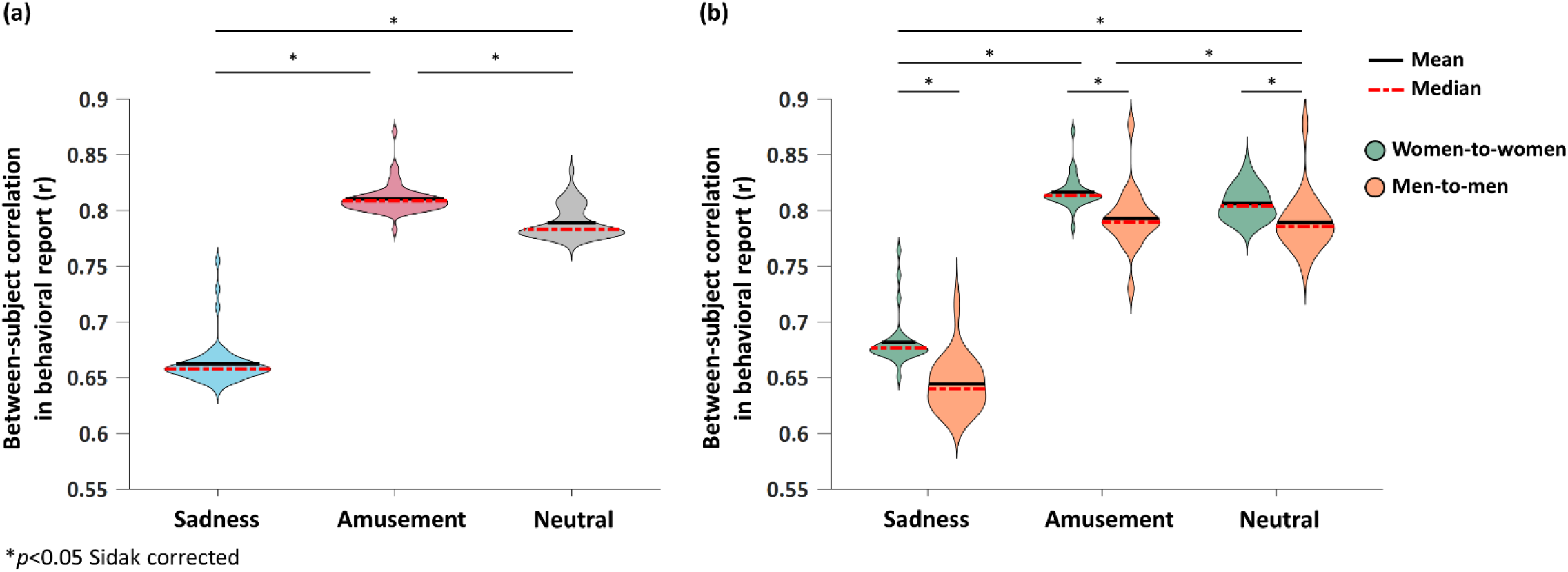
Higher similarity in the behavioral report of emotional experience during amusement and among women. **(a)** The between-subject similarity in the subjective report of emotional experience is presented as a function of the emotional state. Sadness is indicated in blue, amusement in red, and neutral in gray. Higher similarity in the reported emotional experience was found during amusement relative to sadness or neutral. **(b)** The effect of sex was examined by computing women-to-women (green) and men- to-men (orange) similarity. Women were more similar to other women than men to other men, regardless of the emotional state. In each violin, the median is indicated by dashed red lines and the mean by solid black lines. Significant differences are marked by asterisks (*), *p*<0.05 Sidak corrected.

### Modular organization of different brain states

The community structure identified for sadness, amusement, neutral, and rest is presented in Figure 4 (see Supplemental Table S4 for the list of node assignments to modules). Note that visual modules are presented but further excluded from segregation and integration analyses. Each brain state was characterized by a specific topological organization. Eight modules were identified for amusement, neutral, and rest, and nine for sadness. Several qualitative differences in the community structure were observed between the four brain states. The salience network was identified as a separate community only in sadness and amusement. For sadness and neutral, the occipital cortex was divided into two networks: primary and secondary visual, as opposed to one visual network in amusement and rest. Separate sensorimotor and auditory communities were identified in sadness and amusement, whereas in neutral and rest these networks were combined into one community. The posterior default mode network emerged as a separate community only in resting-state.

**Figure 4.**
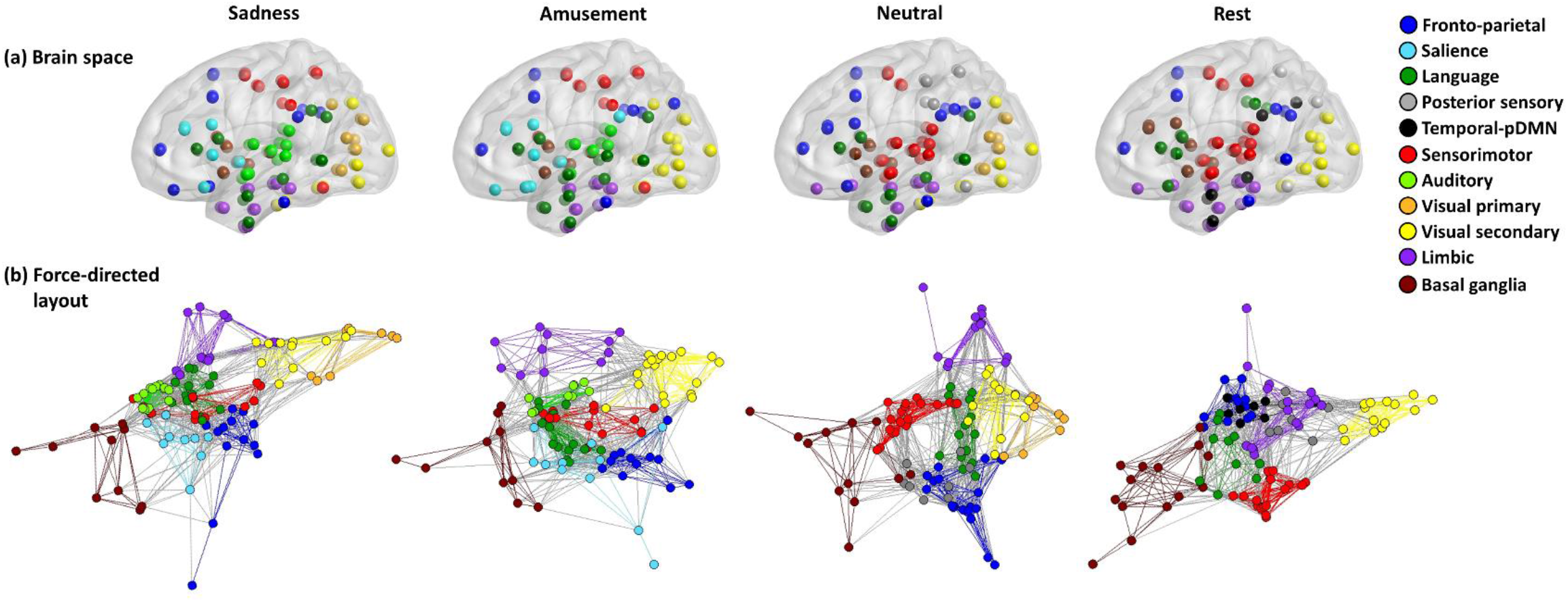
Network communities for different brain states. Communities (i.e., modules) identified for sadness, amusement, neutral, and rest are indicated by colors, and shown in anatomical and topological spaces. **(a)** Anatomical brain space representation of the network communities: brain regions are indicated by circles and colored according to their community assignment. **(b)** Force-directed layouts of the network communities are presented using the Fruchterman-Reingold algorithm (Fruchterman and Reingold 1991). In this layout, connections act as spring-like attractive forces to position nodes in space such that nodes with more shared connections are pulled closer together. The community structure of each brain state was identified using group-level consensus clustering on the individual-level connectivity matrices. For visualization purposes only, the group-averaged functional connections are used here to represent the edges between the nodes in the graphs, and the graphs are displayed at a density of 0.2, i.e., the top 20% of connections are shown for each brain state. Note that visual modules were excluded from segregation and integration analyses. Brain space layouts were visualized using BrainNet Viewer (Xia et al. 2013) and force-directed layouts were visualized using Pajek (Batagelj and Mrvar 1998). pDMN, posterior default mode network.

### Modular segregation and integration during emotional brain states

The mean system segregation (across all modules, excluding visual) was higher for movie states relative to rest (Figures 5a) [main effect of state: *F*(3,147)=11.05, *p*<10^-6^, η_p_^2^=0.18]. The overall mean participation coefficient (across all nodes, excluding visual) was higher for sadness compared to amusement and for both emotional states relative to neutral or rest (Figures 5b) [main effect of state: *F*(3,147)=62.86, *p*<10^-6^, η_p_^2^=0.56]. To compare the segregation and integration of specific modules between the emotional states, further analysis was conducted on the seven modules that were common to both emotions: (1) fronto-parietal; (2) salience; (3) language; (4) sensorimotor; (5) auditory; (6) limbic; and (7) basal ganglia.

**Figure 5.**
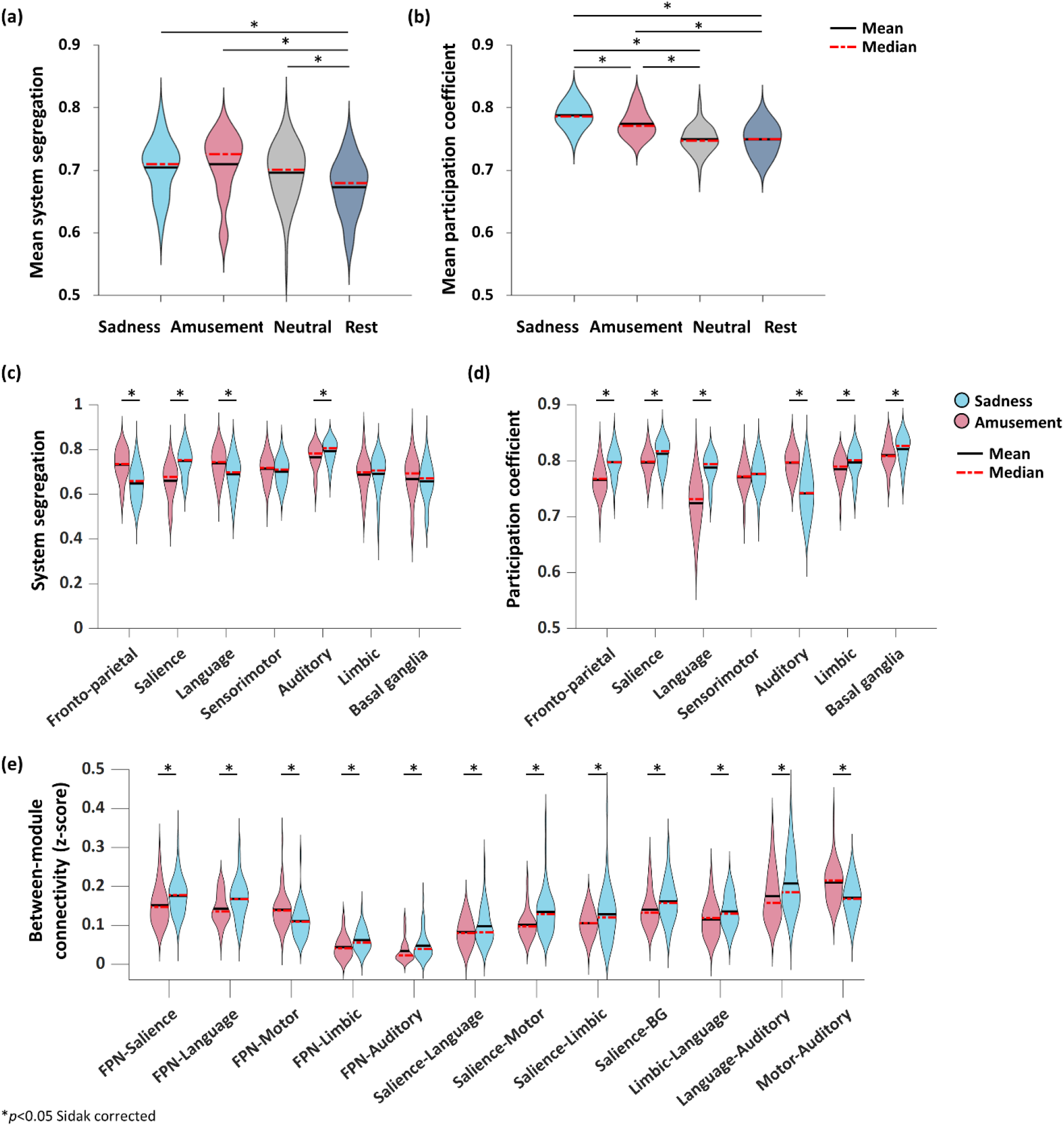
Modular segregation and integration of emotional brain states. **(a)** The mean system segregation (across all modules, excluding visual) is presented as a function of the mental state. Higher values indicate higher segregation. **(b)** The mean participation coefficient (across all nodes, excluding visual) is presented as a function of the mental state. Higher values indicate greater diversity of connections across modules, i.e., higher integration. For sadness (blue) and amusement (red), the **(c)** system segregation, **(d)** participation coefficient, and **(e)** pairwise between-module connectivity are presented as a function of the brain module. Note that Figure 5e presents only the connections that differed between emotional states. In each violin, the median is indicated by dashed red lines and the mean by solid black lines. Significant differences between emotional states are marked by asterisks (*), *p*<0.05 Sidak corrected. BG, basal ganglia; FPN, fronto-parietal network.

For the system segregation, a main effect of module was found (Figure 5c) [*F*(4.92,241.11)=22.29, *p*<10^-6^, η_p_^2^=0.31, Greenhouse-Geisser corrected]. The highest modular segregation was indicated for the auditory module and the lowest segregation for the basal ganglia module. A module-by-emotion interaction was also found [*F*(4.58,224.64)=39.50, *p*<10^-6^, η_p_^2^=0.44, Greenhouse-Geisser corrected]: in sadness relative to amusement, higher segregation was indicated for the salience and auditory modules whereas in amusement higher segregation was indicated for the fronto-parietal and language modules.

For the participation coefficient, main effects of emotion, module, and emotion-by-module interaction were found (Figure 5d) [emotion: *F*(1,49)=21.83, *p*=2.4·10^-5^, η_p_^2^=0.30; module: *F*(4.61,226.13)=65.22, *p*<10^-6^, η_p_^2^=0.57, Greenhouse-Geisser corrected; emotion-by-module: *F*(4.69,229.87)=97.37, *p*<10^-6^, η_p_^2^=0.66, Greenhouse-Geisser corrected]. Across modules, greater diversity of connections, i.e., integration, was indicated in sadness. Furthermore, for both emotions, the highest diversity of connections was indicated for the basal ganglia module. In sadness relative to amusement, higher integration was indicated for the fronto-parietal, salience, language, limbic, and basal ganglia modules whereas in amusement higher integration was indicated for the auditory module.

Specific between-module connections were further examined with pairwise between-module functional connectivity. Main effects of emotion, connection, and emotion-by-connection interaction were found (Figure 5e) [emotion: *F*(1,49)=4.27, *p*=0.044, η_p_^2^=0.08; connection: *F*(8.99,440.62)=97.39, *p*<10^-6^, η_p_^2^=0.66, Greenhouse-Geisser corrected; emotion-by-connection: *F*(9.62,471.53)=7.49, *p*<10^-6^, η_p_^2^=0.13, Greenhouse-Geisser corrected]. Overall greater between-module connectivity was found for sadness, as indicated above by the participation coefficient. In sadness compared to amusement, higher between-module connectivity was found for the frontoparietal-salience, frontoparietal-language, frontoparietal-limbic, frontoparietal-auditory, salience-language, salience-sensorimotor, salience-limbic, salience-basal ganglia, limbic-language, and language-auditory connectivity. In amusement compared to sadness, higher between-module connectivity was found for the frontoparietal-sensorimotor and sensorimotor-auditory connections.

### Relationship between modular metrics and report of emotional experience

During sadness, the mean participation coefficient of the limbic module was associated with the reported valence (*p*=0.008, r=-0.36) and with the intensity of sadness (*p*=0.043, r=0.28). Namely, higher sadness and lower valence were associated with an increased diversity of the limbic module’s connections.

The pairwise between-module connectivity was associated during sadness with the reported valence (frontoparietal-limbic: *p*=1.08·10^-4^, r=-0.52; frontoparietal-sensorimotor: *p*=0.021, r=-0.32; salience-limbic: *p*=0.011, r=-0.35; basal ganglia-limbic: *p*=0.041, r=-0.29; basal ganglia-sensorimotor: *p*=0.034, r=0.29). The intensity of sadness was associated with the connectivity between the frontoparietal-limbic (*p*=0.040, r=0.29) and the salience-limbic (*p*=0.032, r=0.30) modules. The reported arousal during sadness was associated with the connectivity between the salience-limbic modules (*p*=0.029, r=0.34). The results are in line with the higher frontoparietal-limbic and salience-limbic connectivity found in sadness. No associations were found for amusement or with the system segregation.

## Discussion

This study investigated the brain’s network structure during emotional experiences. By using a network science approach, we showed that the segregation and integration of brain network communities are linked to the subjective experience of emotion and reconfigure according to the emotional state. Specifically, we found that sadness was characterized by increased modular integration compared to amusement, measured by a greater diversity of functional connections across modules. Furthermore, the two emotions differed in their between-module connectivity patterns, primarily for the fronto-parietal and salience networks. The fronto-parietal and salience networks are both implicated in task-related activity and cognitive control (Power and Petersen 2013). The salience network is centered on the anterior insula and anterior cingulate cortex and integrates external stimuli with internal states (Seeley et al. 2007). It was shown to mediate the interactions between externally oriented networks, such as the fronto-parietal, and internally oriented networks, particularly the default mode network (Sridharan et al. 2008). The fronto-parietal network includes the dorsolateral prefrontal cortex and the posterior parietal cortex and was associated with working memory and goal-directed behavior (Seeley et al. 2007).

The connections of the fronto-parietal and salience modules, as well as the connectivity between these modules, were stronger in sadness. In addition, the salience module was more segregated in sadness, whereas the fronto-parietal module was more segregated in amusement. Taken together, we found that in sadness compared to amusement, the salience network was both more segregated, i.e., strongly intra-connected, and more integrated, i.e., diversely inter-connected. These metrics are not mutually exclusive, i.e., increased modular segregation does not necessarily imply isolation of the functional community. For example, a previous study showed that modular segregation and global integration increased simultaneously due to the strengthening of specific “hub” connections (Baum et al. 2017).

The salience network emerged as a separate module only in sadness and amusement. In possible congruence, Wager et al. (Wager et al. 2015) demonstrated that sadness and amusement were grouped together according to their preferential activity within the salience network. The fronto-parietal network, on the other hand, was identified in all four mental states (including the non-emotional ones). The fronto-parietal network was previously found to be most variable across tasks and suggested to flexibly interact with other networks to implement task demands (Cole et al. 2013)

Our results resonate with several theoretical accounts of emotion which emphasize a system-level approach (Wager et al. 2015; Pessoa 2019). In general agreement with those theories, we showed that each emotional state can be described by a whole-brain topological structure that involves all functional brain systems rather than a single system or brain region. Adding to those theoretical foundations, we showed that differences between emotional states can be quantified by the level of segregation and integration of functional brain communities/modules.

We showed that the integration between brain modules is related to the behavioral report of emotional valence and intensity. This brain-behavior correspondence demonstrates that the community structure captures important aspects of the subjective emotional experience. These findings may further aid in clinical and behavioral prediction. Behavioral prediction was argued to be more accurate when brain measures are taken from a relevant cognitive state (Greene et al. 2018). Thus, characterizing the brain’s community structure during ecological states of sadness and amusement (and other emotional states such as fear or stress, for example) may enhance our ability to predict prospective psychiatric symptoms in at-risk and clinical populations.

Contrary to our hypothesis, the emotional condition did not consistently increase the within- and between-subject similarity in the functional connectome, but rather this effect was dependent on the brain parcellation atlas. With respect to the within-subject similarity, an individual’s functional connectome was more similar between two emotional states, only for the HO and Schaefer atlases. An even greater effect of the brain parcellation atlas was indicated for the between-subject similarity: for the HO and Shen atlases, movie-watching increased the similarity between individuals relative to rest, whereas an opposite effect was found for the AAL and Schaefer atlases. This result may explain some previous conflicting findings, where several studies showed that movies increase synchronization between cortical signal fluctuations across individuals (Hasson et al. 2004; Lankinen et al. 2014) while others found reduced between-subject similarity during movie-watching relative to rest (Geerligs et al. 2015). We note that in contrast to previous work, we excluded the entire occipital cortex from our similarity analyses, to reduce the impact of visual properties of the stimuli.

Furthermore, we provided a quantitative characterization of differences in modular organization between movie states and rest. We found that movie-watching, compared to rest, was characterized by higher modular segregation. This is in agreement with a recent study (Kim et al. 2018) which indicated reorganization of the community structure during movie-watching relative to rest, however, it did not quantify the modular segregation or integration of the states. We note that studies that quantitatively compared functional connectivity between different movies are scarce, and most studies that examined task or state effects on individual differences in the functional connectome did not include naturalistic stimuli (Cole et al. 2014; Gratton et al. 2018).

The similarity between individuals in the functional connectome was reduced with the use of smaller parcels. One possible explanation is that smaller parcels are more sensitive to anatomical variability and imperfect coregistration, and to functional variability, namely, slightly different anatomical locations for the same function across participants. Furthermore, Salehi et al. (Salehi et al. 2020) showed that brain subdivisions based on functional connectivity data differ according to the task during which the data was acquired, even within an individual. In other words, parcels that were defined based on resting-state data, may not capture accurate functional units during other tasks or mental states.

The magnitude of sex differences in functional connectome similarity across individuals varied according to the mental state, in agreement with recent findings (Finn et al. 2017). Notably, Finn and colleagues found the largest sex differences during an emotion perception task. In a related study, Green et al. (Greene et al. 2018) showed that the task that increases the ability to predict fluid intelligence from functional connectomes differed by sex.

In the behavioral reports of emotion, we found an overall higher similarity among women. In addition, amusement yielded the highest similarity in behavioral ratings among participants, compared to sadness or the neutral state. Emotion similarity in the subjective experience among individuals is known to be associated with many interpersonal advantages, such as greater satisfaction, empathy, cooperation, and reduced stress (Locke and Horowitz 1990; Barsade 2002; Townsend et al. 2014). Our results suggest that similarity in the experienced emotion, and thus its positive social effects, are more likely to be achieved during states of amusement and among women.

## Limitations

Emotional states were elicited by movies, using different movie clips for each state (sadness, amusement, neutral). To minimize the impact of visual properties of the movie stimuli on brain states, we have excluded the occipital cortex from all analyses. In addition, our analyses indicated no differences between states in low- and mid-level visual or auditory features of the movie stimuli. However, it is important to note that high-level visual information (e.g., scenes, objects, actions) which is processed outside the visual cortex, can contribute to differences between brain states. This includes inferior temporal regions within the ventral visual stream that respond to objects, scenes, and faces (among other stimuli), as well as parietal and frontal regions within the dorsal stream or the dorsal attention network, which respond to a wide range of visual features, including object location (Corbetta and Shulman 2002). In addition, the ventral posterior medial network, i.e., the parahippocampal cortex, retrosplenial cortex, and posterior angular gyrus, was shown to respond to event transitions during movie-watching (Cooper et al. 2021). While the possible contribution of high-level non-emotional features cannot be ruled out, the following was done to minimize their influence. First, our analyses took a static functional connectivity approach and not a dynamic one. Namely, correlations were computed between time series across the entire length of acquisition and not across short timescales within the data. Second, each emotional state was elicited by four movie clips, thus the state-specific functional connectomes captured the average interactions between brain regions, across the different movies that induced a certain state. Characterizing states by averaging over several different movies instead of one was chosen to increase the generalizability of the findings and reduce the impact of movie-specific features, in addition to allowing for a continuous, naturalistic, and intense emotional experience.

Modularity analyses were conducted using the HO parcellation atlas. As indicated by the between-subject functional connectivity similarity analysis, the choice of the parcellation atlas may impact the results. Studies have started to examine the influence of the parcellation atlas on the reliability of graph theoretical metrics (Bertolero et al. 2015; Cao et al. 2019; Tozzi et al. 2020), yet more work is needed to understand its impact, and in particular on metrics of modular segregation and integration. We also note that while the order of the emotional states (sadness, amusement) was balanced across the sample and within each sex, the order of the rest and neutral states was not. Rest was always the first state, and neutral was induced between the emotional states. Thus, possible effects of fatigue can influence the comparison between the states.

Last, the study design does not enable us to conclude whether the findings are specific to the emotional categories (sadness, amusement) or emotional valence (negative, positive). Moreover, despite the discreteness of the rating results for the target emotions, the mapping between emotional states and modular brain representations can be of many-to-one (and not one-to-one). In other words, the modular representation of anger, for example, can be similar to that of sadness. We note that recent studies did not provide strong support for traditional dimensional approaches to emotion at the level of the brain’s network organization (Kragel and LaBar 2015; Wager et al. 2015). Specifically, emotional categories were not indicated to be grouped according to their valence. For example, sadness and amusement, despite their contrasting valence, were more similar to one another in their brain representations than emotions of the same valence, such as sadness and fear (Wager et al. 2015). In another study, neural activity patterns were most separable during the experience of distinct emotional categories and not differentiable according to valence and arousal (Kragel and LaBar 2015). Studies examining the functional connectome similarity and modular organization during additional emotional and affective states are needed.

## Conclusions

By applying network science methods to the neural representations of intense naturalistic emotional states, this study sought to deepen our understanding of the brain basis of emotion in humans. A modular organization of the brain during experiences of sadness and amusement was described, and the results extended previous attests for the essential importance of the brain’s network segregation and integration patterns, to the field of emotion. Our results suggest that the experience of emotion is linked to a reconfiguration of distributed, not emotion-specific, functional brain networks. The interaction patterns between functional networks, and not the networks themselves, are postulated to be associated with the emotional state.

## CRediT authors statement

**Rotem Dan:** Conceptualization; Data curation; Formal analysis; Investigation; Project administration; Resources; Visualization; Writing - Original draft; Writing – Review & Editing. **Marta Weinstock:** Resources; Supervision; Writing – Review & Editing. **Gadi Goelman:** Resources; Supervision; Writing – Review & Editing.

## Code availability

The Brain Connectivity Toolbox (BCT) was used for modularity analyses and is freely available at https://sites.google.com/site/bctnet. Consensus partitioning was done using the following BCT functions: community_louvain.m, agreement.m, consensus_und.m. The weighted participation coefficient was calculated using the participation_coef.m function from the BCT. The normalized mutual information (NMI) was computed using getNMI.m function and is available at https://www.mathworks.com/matlabcentral/fileexchange/62974-getnmi-a-b.

