## Supplemental Material for "Emotional states as distinct configurations of functional brain networks"

### Supplemental Methods

- Movie clips used for the induction of emotional states
    - Table S1. Description of movie stimuli.
    - Movie rating questionnaire collected outside the scanner.
- MRI data acquisition
- MRI data preprocessing
- Choice of the resolution parameter and treatment of negative weights in Louvain modularity
    - Figure S1. Between-subject similarity in the community structure as a function of gamma and the treatment of negative weights.
    - Figure S2. The number of individual-level modules as a function of gamma and the treatment of negative weights.
    - Figure S3a. Average similarity between community structures of different resolutions as a function of gamma and the treatment of negative weights.
    - Figure S3b. The similarity between community structures of different resolutions for a modularity function that includes only positive weights.
    - Figure S4. The stability between runs of group-level community identification as a function of gamma and the treatment of negative weights.
    - Figure S5. The number of group-level modules as a function of gamma and the treatment of negative weights.

### Supplemental Results

- Table S2. Visual and auditory features of movie stimuli.
- Table S3. Discreteness analysis of the target emotional state.

- Figure S6. Within-subject similarity in the functional connectome is not dependent on sex.
- Figure S7. Within-subject similarity in the functional connectome is not dependent on sex: analysis of a balanced subsample of 20 women and 20 men.
- Figure S8. Between-subject similarity in the functional connectome: analysis of a balanced subsample of 20 women and 20 men.
- Figure S9. Between-subject similarity in the behavioral report: analysis of a balanced subsample of 20 women and 20 men.
- Table S4. The community structure identified for sadness, amusement, neutral, and rest.

### Supplemental Methods

#### Movie clips used for the induction of emotional states

**Table S1. Description of movie stimuli.**

| Movie | Content | Target emotion | Clip length (min:sec) | Set of stimuli |
|---|---|---|---|---|
| Stepmom | Jackie tells her children that she is dying of cancer, resulting in her daughter emotionally storming out | sadness | 3:01 | Gross and Levenson 1995 |
| Lion King | Lion cub finds his dead father in a gorge | sadness | 2:11 | Rottenberg et al. 2007 |
| Terms of Endearment | Dying Emma says goodbye to her children before her death in the hospital | sadness | 3:02 | Farb et al. 2010 |
| The Champ | Boy witnesses his father dying after a boxing match | sadness | 3:01 | Gross and Levenson 1995 |
| There's Something About Mary | Ted fights with a dog | amusement | 3:02 | Schaefer et al. 2010 |
| Benny and Joone | Benny plays the fool in a coffee shop | amusement | 2:10 | Schaefer et al. 2010 |
| Bill Cosby | Bill Cosby does stand-up comedy | amusement | 2:02 | Rottenberg et al. 2007 |
| When Harry met Sally | Sally simulates an orgasm in a restaurant | amusement | 2:51 | Rottenberg et al. 2007 |

The induction of amusement was composed of the four movie clips presented in a row and lasted 10 minutes and 5 seconds. The induction of sadness was composed of the four movie clips presented in a row and lasted 11 minutes and 15 seconds. For both mood inductions, only the first 10 minutes were used for the functional connectivity MRI analysis. Neutral movie clips were taken from television programs on gardening, painting, jewelry making, and cooking.

**Movie rating questionnaire**

**Block: ___**

The following questions refer to how you felt *while watching the movies.*

| 1 | 2 | 3 | 4 | 5 | 6 | 7 | 8 |
|---|---|---|---|---|---|---|---|
| Not at all | | | Somewhat | | | | Extremely |

Using the scale above (1-8), please indicate the amount of each emotion you experienced while watching the movie clips.

| | | |
|---|---|---|
| ____________Disgust | ____________Enjoyment | ____________Embarrassment |
| ____________Amusement | ____________Sadness | ____________Peacefulness |
| ____________Anger | ____________Interest | ____________Confusion |
| ____________Sorrow | ____________Contempt | ____________Fear |
| ____________Guilt | ____________Happiness | ____________Surprise |
| ____________Anxiety | ____________Calm | ____________Shame |

Did you feel any other emotion during the movies? ◯ No ◯ Yes

If so, what was the emotion?_______________________

How much of this emotion did you feel? (use the scale above) _______________________

Use the following scale to indicate your general feeling of pleasantness, from pleasant/positive to unpleasant/negative, while watching the movies. Please circle your answer:

1 2 3 4 5 6 7 8

Negative Positive

Unpleasant Pleasant

Use the following scale to indicate your arousal while watching the movies. Please circle your answer:

1 2 3 4 5 6 7 8

Low High

Calm Excited/alerted

Have you seen any of the movies before? ◯ No ◯ Yes

Did you fall asleep during any part of the movies? ◯ No ◯ Yes

### MRI data acquisition

Functional images were acquired using $T_2^*$-weighted gradient-echo echo-planar imaging (GE-EPI) sequence with TR=2 sec, TE=30 ms, image matrix=64×64, field of view=192x192 mm, flip angle=90°, resolution=3x3x3 mm, interslice gap=0.45 mm. Each brain volume comprised 30 axial slices. Anatomical images were acquired using a sagittal T1-weighted MP-RAGE sequence with TR=2.2 sec, TE=2.43 ms, resolution=1x1x1 mm. The T1-weighted images were acquired for the coregistration and normalization of the functional images.

### MRI data preprocessing

Preprocessing of fMRI data was done using SPM12 software (Wellcome Trust Centre for Neuroimaging, London, United Kingdom, http://www.fil.ion.ucl.ac.uk/spm/software/spm12). Functional images were spatially realigned and unwarped, slice-timing corrected, segmented, and normalized to MNI space. Further preprocessing was done in CONN toolbox (Whitfield-Gabrieli and Nieto-Castanon 2012). Potential confounding effects were regressed out using the aCompCor method for anatomical component-based noise correction (Behzadi et al. 2007). These included: (i) outlier scans, removed by censoring/scrubbing (Power et al. 2014). Outlier scans were identified based on the amount of subject motion in the scanner as measured by the framewise displacement (FD) and global BOLD signal. Acquisitions with FD > 0.9mm or global BOLD signal changes > 5 standard deviations were considered outliers and removed by regression. (ii) First 5 principal components (PCAs) of the CSF and white matter signals: were regressed out to minimize the effects of physiological non-neuronal signals such as cardiac and respiratory signals. Prior to the regression of PCAs, the white matter and CSF masks were eroded, i.e., only voxels with values above 50% in the white matter and CSF posterior probability maps were used, to ensure the regression of pure white matter or CSF signals. (iii) Estimated subject-motion parameters and their first-order derivatives (a total of 12 parameters). (iv) Session effects: The potential effects of the beginning of the session were removed by a step function convolved with the hemodynamic response function, in addition to the linear BOLD signal trend. Note that global signal regression was not performed. After regression of all potential confounding effects, temporal band-pass filtering (0.008-0.09 Hz) was performed.

One participant was excluded due to more than 15% censored TRs per state (specifically, 16% of TRs for amusement). For the remaining sample, the average percentage of data removed by censoring per state per participant was: (i) sadness: 1.28±1.88% (range: 0-9%); (ii) amusement: 2.26±2.89% (range: 0-12.3%); (iii) neutral: 1.61±2.04% (range: 0-8.6%); (iv) rest: 0.71±1.36% (range: 0-6%).

To quantify state and sex differences in head motion, we calculated the mean framewise displacement (FD) and the root mean square displacement (RMS) using the approach of Power et al. (2014). Differences in motion measures (mean FD, RMS) were tested by two-way mixed ANOVA models implemented by a GLM with the state (sadness, amusement, neutral, rest) as a within-subject factor and sex (women, men) as a between-subject factor. The models tested for: (i) main effect of state; (ii) main effect of sex; (iii) sex-by-state interaction. There were no significant effects.

**Choice of the resolution parameter and treatment of negative weights in Louvain modularity**

Consensus partitioning was conducted for each state at the individual and group levels over a range of the resolution parameter gamma (Reichardt and Bornholdt 2006): from 1 to 3, in increments of 0.25. The resolution parameter impacts the number of identified communities within a network: small values of gamma yield fewer communities and larger values enable to detect more communities that are smaller in size.

In addition, we examined three variants of the weighted modularity function (Q) (Rubinov and Sporns 2011): positive only, symmetric treatment of positive and negative weights, and asymmetric treatment of negative weights. The origin and interpretation of negative functional correlations remain unclear (Fox et al. 2009). Recent developments were made to incorporate both positive and negative weights in the modularity function (Rubinov and Sporns 2011). While positive weights indicate common activity patterns and thus support the placement of nodes within the same module, negative weights are thought to indicate anti-phase or decoupled activity and thus support the placement of nodes in distinct modules.

For each value of gamma, we tested the following three variants of the weighted modularity function (Rubinov and Sporns 2011): (i) positive only: setting the negative weights to zero and maximizing positive within-module connections; (ii) symmetric: equal weight on maximizing positive within-module connections and minimizing negative within-module connections; (iii) asymmetric: greater weight on maximizing positive within-module connections. In the asymmetric function, the contribution of negative weights decreases with an increase in positive weights. For all runs of the Louvain modularity algorithm, a threshold of tau=0.05 was used to set cells with an agreement of less than tau to zero (Lancichinetti and Fortunato 2012).

To compute the similarity in community structure between participants, states, gamma values, or runs, we used the normalized mutual information (NMI) (Kuncheva and Hadjitodorov, 2004). The NMI measures the dependency between two node assignments, i.e., how much information one set of assignments provides about a second set of assignments, and ranges from 0 (no information/similarity) to 1 (identical community structure).

We chose a resolution parameter of gamma=2.25 and a positive-weighted modularity function (without negative connections) based on the following considerations, at the individual and group levels:

*At the individual level:*

(i) High between-subject similarity in the community structure per state (Supplementary Figure S1). A higher between-subject similarity was found for the positive-weighted modularity function, compared to the symmetric and asymmetric functions.

(ii) Minimal number of singleton modules (i.e., communities that include one node) (Supplementary Figure S2). For gamma approximately greater than 2.5 for sadness and amusement, and gamma greater than 2.25 for neutral and rest, many singleton modules were identified using the symmetric and asymmetric modularity functions.

Note that there is a tradeoff between criteria (i) and (ii) with respect to gamma: increasing gamma increases the between-subject similarity in the community structure but also increases the identification of singleton modules.

*At the group level:*

(iii) High similarity of the community structure to partitions obtained with other gamma values (Supplementary Figures S3a,b). For the positive-weighted modularity function, gamma=2.25 was the minimal gamma value included in the ranges of partition stability for all states.

(iv) Stability/ low degeneracy of the identified group community structure across different runs at the group-level. To validate the stability of the identified group-level community structure, we repeated for each state the group-level consensus partitioning 50 times and examined the similarity between runs (Supplementary Figure S4). For the positive-weighted function, gamma=2.25 resulted in no degeneracy for amusement, neutral and rest, and low degeneracy for sadness.

(v) Number of communities identified at the group level which is comparable to that identified at the subject-level (Supplementary Figure S5). This criterion is related to criterion (ii) of minimizing singleton communities at the individual level, resulting in comparable numbers of communities at the individual and group levels.

**Figure S1. Between-subject similarity in the community structure as a function of gamma and the treatment of negative weights**

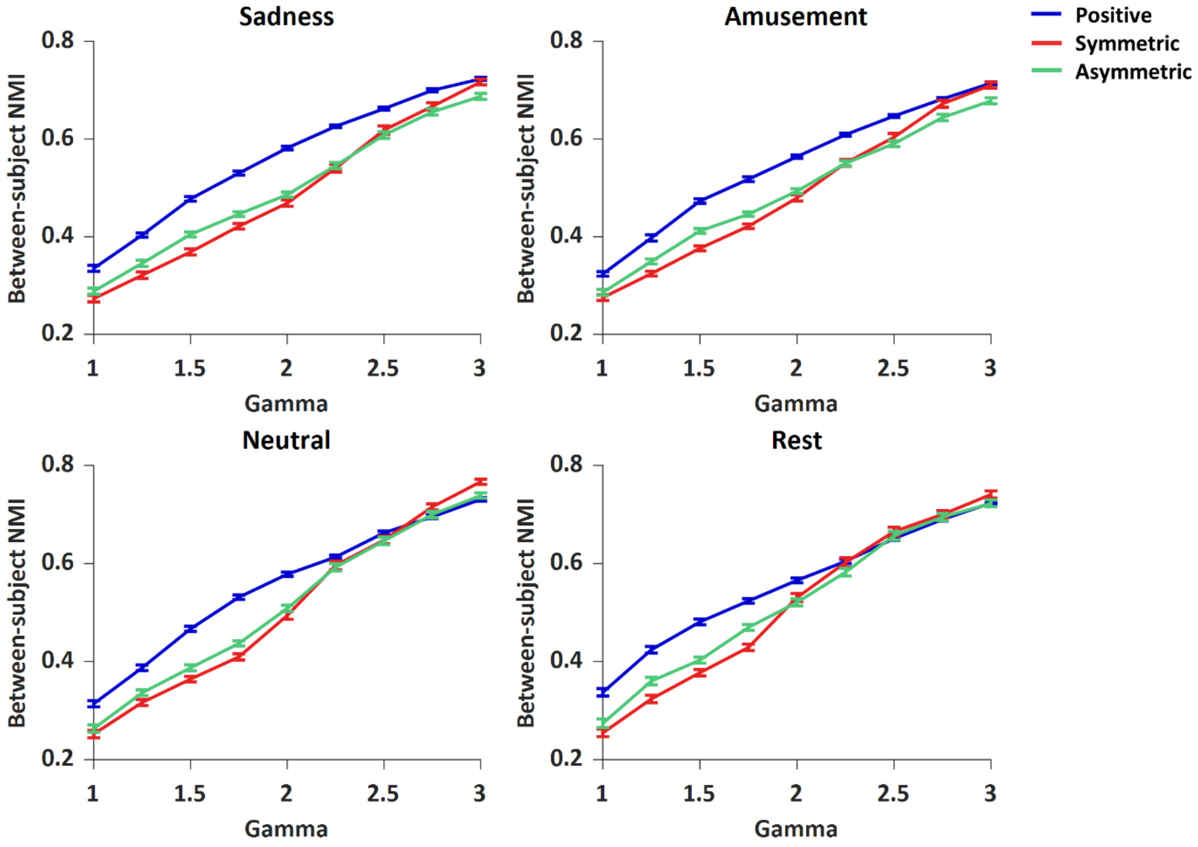


The similarity between the community structure of different participants was measured by the NMI and shown on the y-axis, different values of the resolution parameter (gamma) are indicated on the x-axis, and variants of the modularity function are indicated by colors. Average values are shown with standard errors. Increasing the resolution parameter increased the between-subject similarity. A higher between-subject similarity was found for the positive-weighted modularity function, compared to the symmetric and asymmetric functions. Note that as shown in Figure S2, for high values of gamma, many singleton communities (i.e., including one node) were present, especially when using the symmetric and asymmetric modularity functions. These singleton communities introduced artificial high between-subject similarity such that it reached that of the positive-weighted modularity function.

**Figure S2. Number of individual-level modules as a function of gamma and the treatment of negative weights**

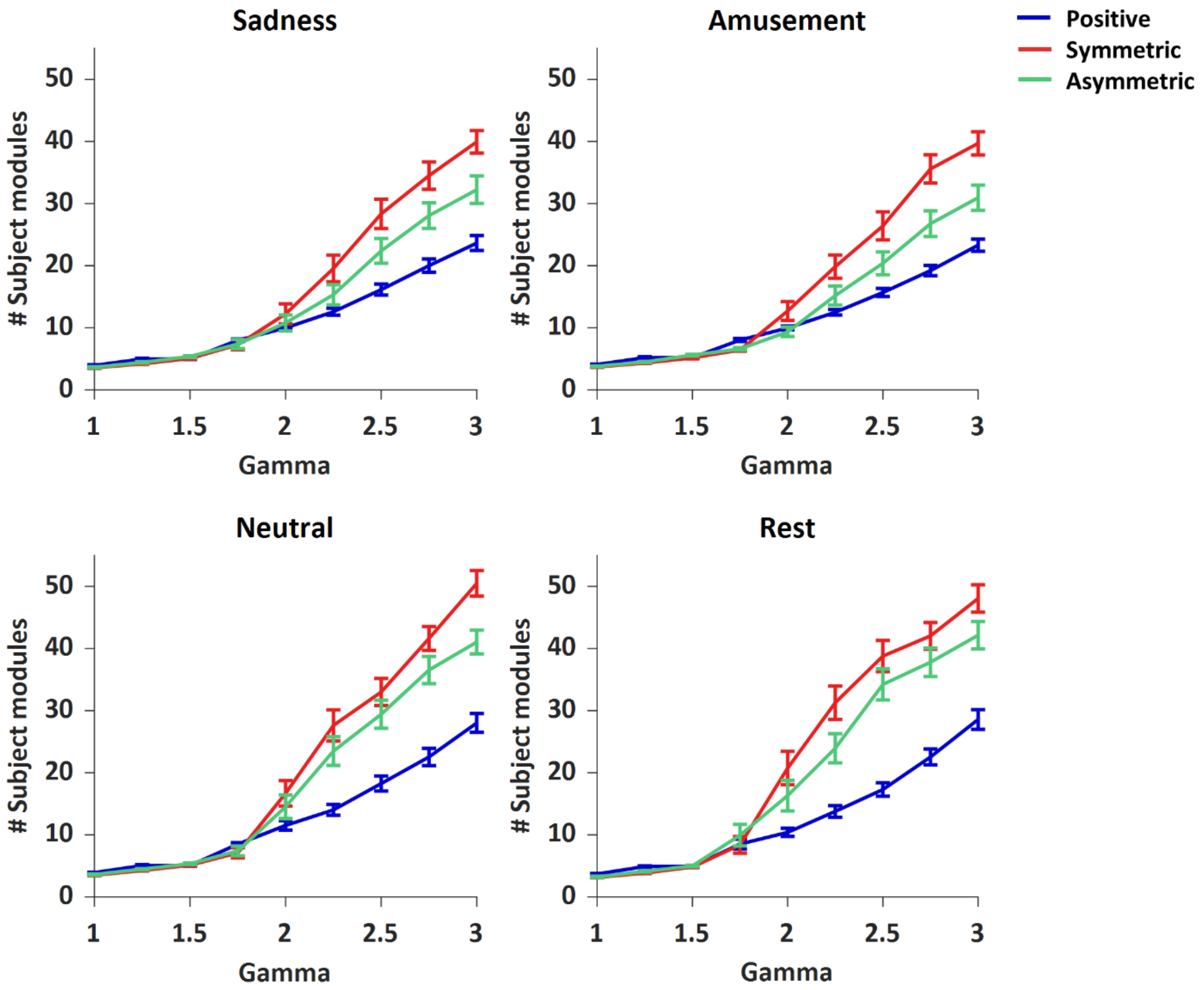


The number of modules identified for a single subject is shown on the y-axis, different values of the resolution parameter (gamma) are indicated on the x-axis, and variants of the modularity function are indicated by colors. Average values are shown with standard errors. The number of modules increased with the resolution parameter gamma. After a certain value of gamma (~2.5 for sadness and amusement, ~2.25 for neutral and rest) many singleton modules (i.e., including one node) were identified when using the symmetric and asymmetric modularity functions.

**Figure S3a. Average similarity between community structures of different resolutions as a function of gamma and the treatment of negative weights**

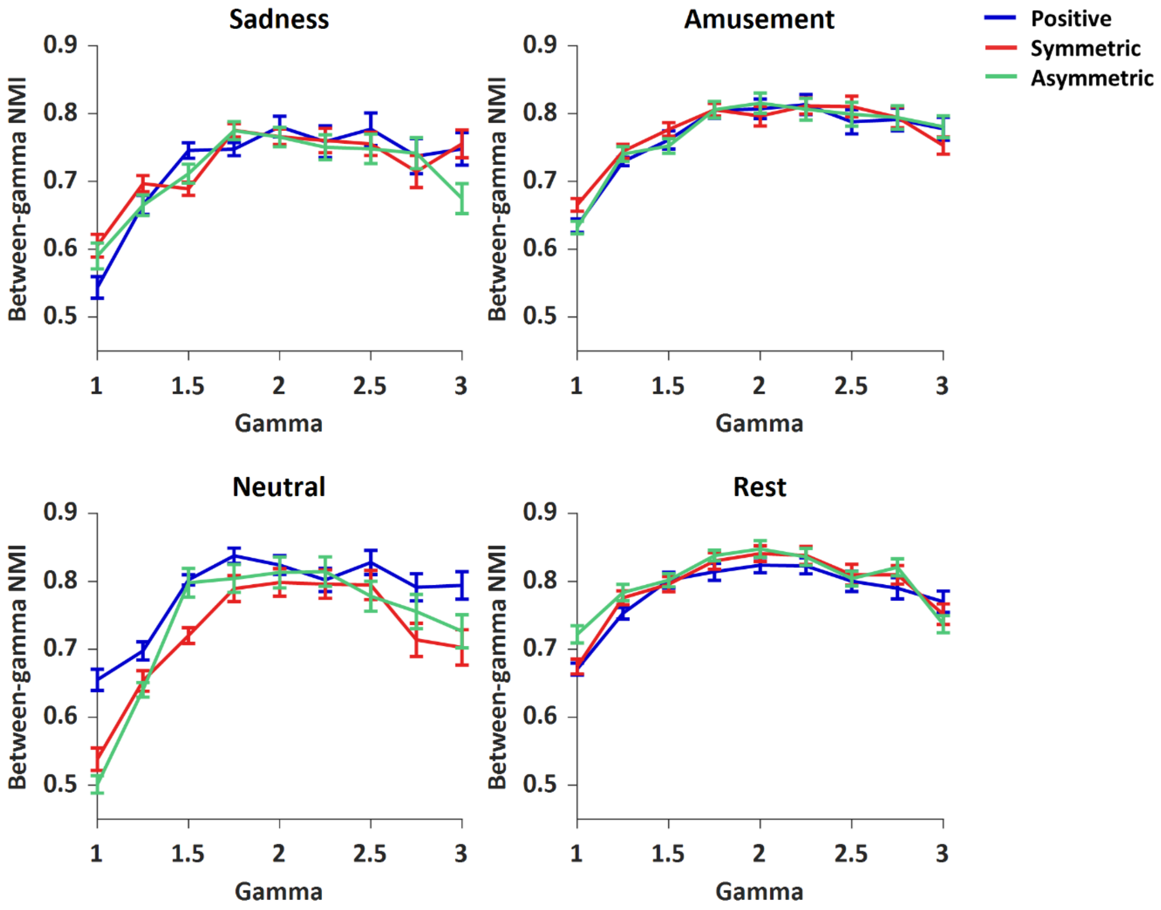


The average similarity between the community structures obtained with different gammas was computed by the NMI and is shown on the y-axis, different values of the resolution parameter (gamma) are indicated on the x-axis, and variants of the modularity function are indicated by colors. Average values are shown with standard errors. For the positive-weighted modularity function, the similarity in community structure between pairs of gamma values is shown in Figure S3b.

**Figure S3b. Similarity between community structures of different resolutions for a modularity function that includes only positive weights**

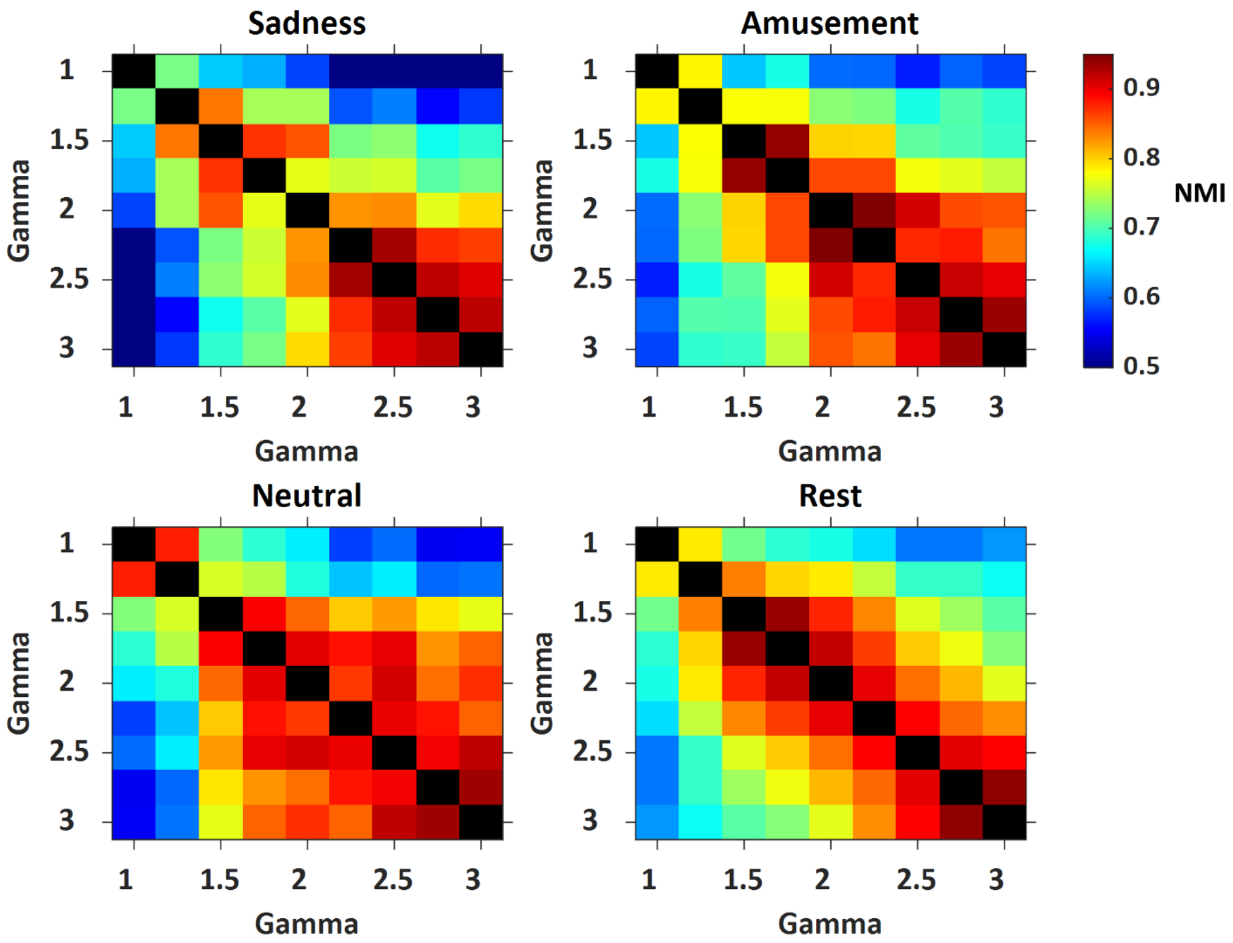


For the positive-weighted modularity function, each cell in the matrices shows the level of similarity (measured by NMI) between a pair of gamma values, as indicated by the color bar. The following ranges of gamma values were found to show high similarity, suggesting stabile partitioning:

Sadness: [2.25 3]; amusement: [2 3]; neutral: [1.75 3]; rest: [1.5 2.25] and [2.25 3].

Note that gamma=2.25 is the minimal gamma value that is included in the above ranges of stability for all states.

**Figure S4. Stability between runs of group-level community identification as a function of gamma and the treatment of negative weights**

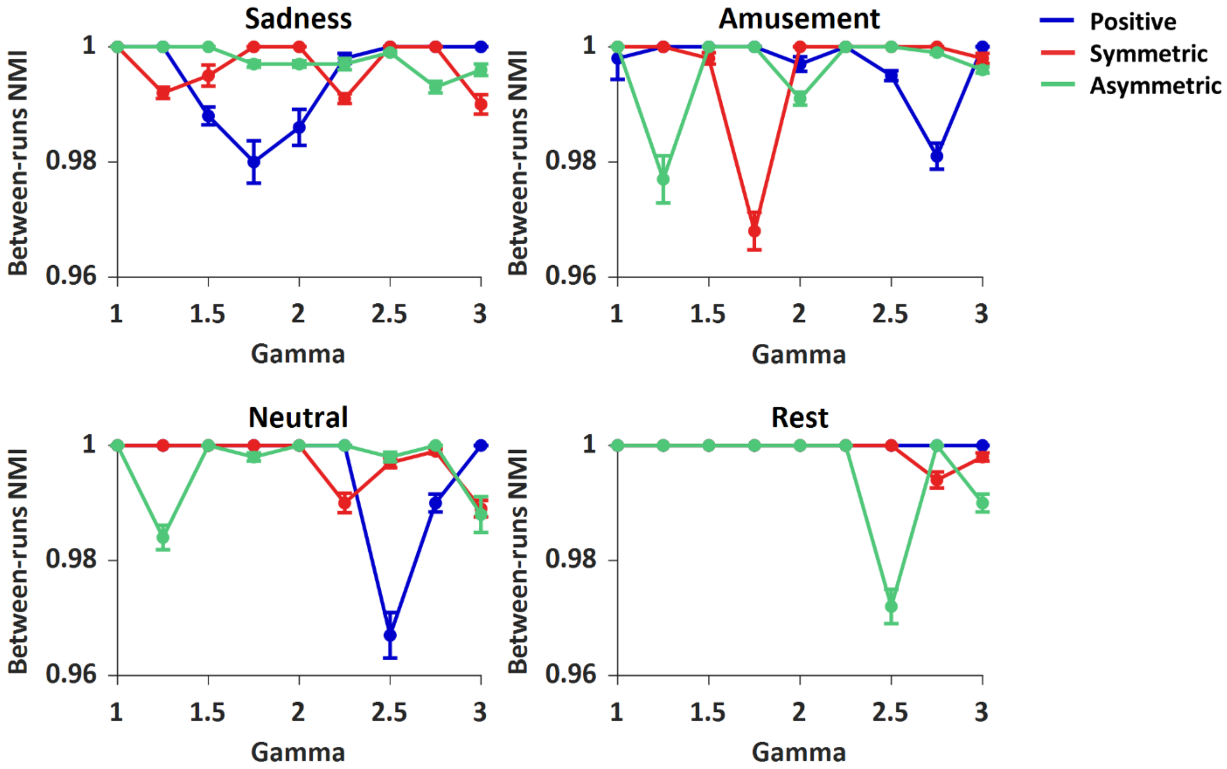


The similarity between different runs of the group-level community detection was measured by the NMI and is shown on the y-axis, different values of the resolution parameter (gamma) are indicated on the x-axis, and variants of the modularity function are indicated by colors. Average values are shown with standard errors. An NMI value less than 1 indicates degeneracy, namely more than one near-optimal partition that maximized the modularity function. For example, for sadness, for gamma=2.25 and a positive-weighted function, 48/50 runs of the group-level partitioning algorithm yielded one solution for the community structure and 2/50 runs yielded another solution, resulting in an NMI <1 (0.998±0.006). Higher degeneracies were found for sadness for gamma values in the range of [1.5 2]. For amusement, neutral, and rest, only one solution was found for all runs with gamma=2.25 and a positive-weighted function.

**Figure S5. Number of group-level modules as a function of gamma and the treatment of negative weights**

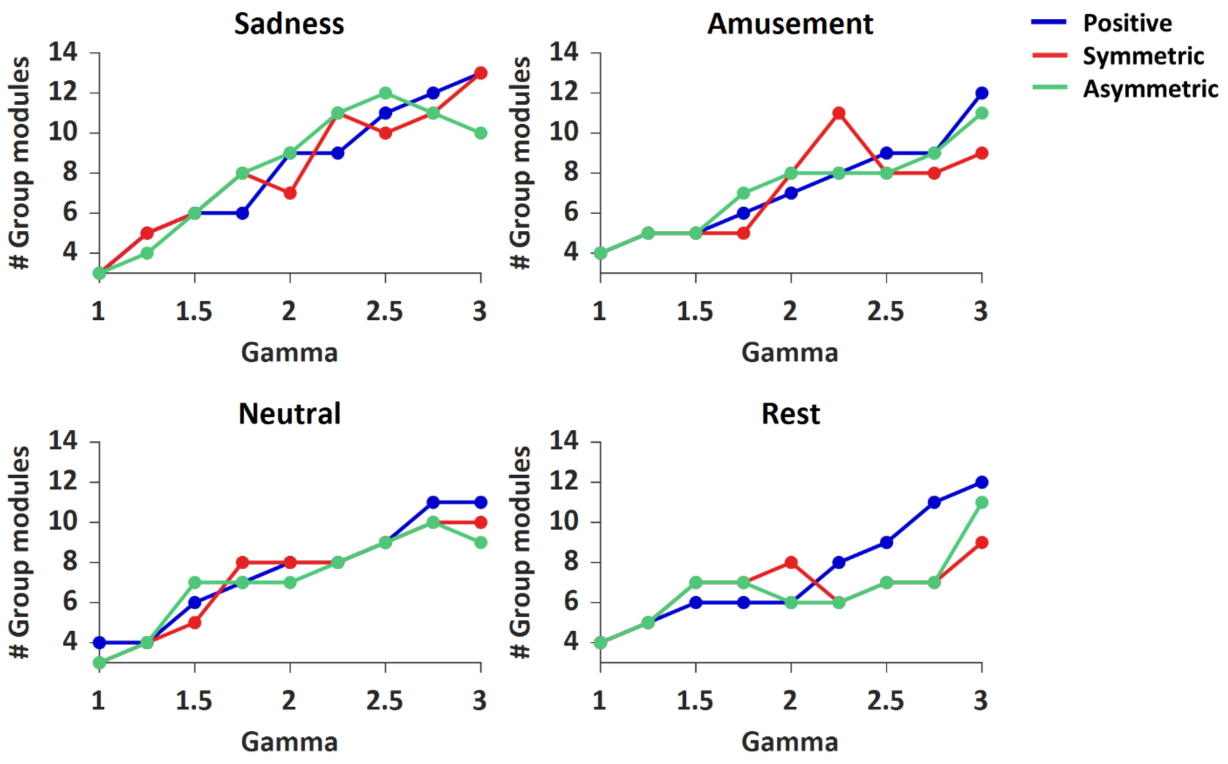


The number of group-level modules is shown on the y-axis, different values of the resolution parameter (gamma) are indicated on the x-axis, and variants of the modularity function are indicated by colors.

### Supplemental Results

**Table S2. Visual and auditory features of movie stimuli.**

| | Brightness (mean±std) | Vibrance (mean±std) | Audio RMSE (mean±std) | Optic flow (mean±std) | Faces⁺, *Pliers* analysis | Faces⁺, manual encoding (mean±std) |
|---|---|---|---|---|---|---|
| **Amusement** | | | | | | |
| Benny and Joone | 0.33±0.02 | 169.01±52.40 | 0.02±0.01 | $9.27 \cdot 10^5 \pm 9.10 \cdot 10^5$ | 0.61 | 1.43±0.80 |
| Bill Cosby | 0.15±0.08 | 244.93±210.29 | 0.08±0.03 | $5.77 \cdot 10^5 \pm \cdot 5.71 \cdot 10^5$ | 0.85 | 1.00±0.00 |
| There's Something About Mary | 0.32±0.05 | 399.12±216.26 | 0.03±0.02 | $2.28 \cdot 10^5 \pm 3.53 \cdot 10^5$ | 0.41 | 0.99±0.78 |
| When Harry met Sally | 0.34±0.04 | 348.76±87.79 | 0.03±0.03 | $3.50 \cdot 10^5 \pm 4.53 \cdot 10^5$ | 0.83 | 1.65±1.30 |
| **Sadness** | | | | | | |
| Stepmom | 0.24±0.07 | 184.00±57.97 | 0.03±0.04 | $1.09 \cdot 10^5 \pm 1.88 \cdot 10^5$ | 0.78 | 1.29±0.70 |
| Lion King | 0.46±0.07 | 828.85±280.85 | 0.02±0.04 | $3.88 \cdot 10^5 \pm 7.21 \cdot 10^5$ | - | 1.04±0.79 |
| Terms of Endearment | 0.42±0.06 | 311.57±121.68 | 0.01±0.01 | $4.48 \cdot 10^5 \pm 6.59 \cdot 10^5$ | 0.69 | 1.28±0.57 |
| The Champ | 0.13±0.03 | 54.32±21.26 | 0.01±0.01 | $7.55 \cdot 10^4 \pm 7.55 \cdot 10^4$ | 0.46 | 1.66±1.06 |
| **Neutral** | | | | | | |
| Cooking | 0.63±0.02 | 1313.86±265.38 | 0.04±0.03 | $1.70 \cdot 10^5 \pm 2.51 \cdot 10^5$ | 0.87 | 1.89±0.40 |
| Gardens | 0.28±0.03 | 550.68±144.74 | 0.02±0.02 | $1.52 \cdot 10^5 \pm 2.48 \cdot 10^5$ | 0.56 | 1.43±0.80 |
| Jewelry | 0.67±0.02 | 102.74±43.62 | 0.03±0.03 | $3.13 \cdot 10^4 \pm 7.36 \cdot 10^4$ | 0.90 | 1.90±0.42 |
| Paint | 0.45±0.05 | 631.63±279.19 | 0.03±0.03 | $3.09 \cdot 10^5 \pm 3.22 \cdot 10^5$ | 0.31 | 0.78±0.94 |

Means and standard deviations were calculated across frames.

⁺Faces: were analyzed both by *Pliers* and by manual encoding. Note that *Pliers* does not detect animated faces, thus it did not detect faces for the Lion King. The following metrics were computed and shown in the table: (i) *Pliers*: the proportion of frames containing at least one face, (ii) manual encoding: the number of faces presented for each frame (means and standard deviations).

There were no statistical differences between states for any of the movie features.

**Table S3. Discreteness analysis of the target emotional state.**

| Discreteness measure | Sadness movie state (target: sadness) | | Amusement movie state (target: amusement) | |
|---|---|---|---|---|
| | Outside scanner | Within scanner | Outside scanner | Within scanner |
| Target vs. fear | $p$=7.2·10$^{-10}$ | $p$=1.3·10$^{-8}$ | $p$=7.0·10$^{-10}$ | $p$=6.6·10$^{-10}$ |
| Target vs. anger | $p$=1.0·10$^{-9}$ | $p$=9.7·10$^{-10}$ | $p$=6.9·10$^{-10}$ | $p$=5.5·10$^{-10}$ |
| Target vs. sadness/ amusement (depending on target) | $p$=1.2·10$^{-9}$ | $p$=4.9·10$^{-8}$ | $p$=6.5·10$^{-10}$ | $p$=5.0·10$^{-10}$ |
| Target vs. embarrassment | $p$=7.1·10$^{-10}$ | - | $p$=8.6·10$^{-10}$ | - |
| Target vs. disgust | $p$=7.1·10$^{-10}$ | - | $p$=7.6·10$^{-10}$ | - |
| Target vs. surprise | $p$=1.0·10$^{-9}$ | - | $p$=1.3·10$^{-8}$ | - |
| Target vs. guilt | $p$=1.0·10$^{-9}$ | - | $p$=6.7·10$^{-10}$ | - |
| Target vs. anxiety | $p$=7.2·10$^{-10}$ | - | $p$=7.3·10$^{-10}$ | - |
| Target vs. contempt | $p$=7.1·10$^{-10}$ | - | $p$=8.6·10$^{-10}$ | - |
| Target vs. confusion | $p$=1.0·10$^{-9}$ | - | $p$=7.0·10$^{-10}$ | - |

For both emotional states, all pairwise comparisons between the intensities of the target emotion and other emotion categories were significant (all $p<1.9\cdot10^{-3}$, Bonferroni correction), indicating discrete emotional experiences of sadness and amusement.

**Figure S6. Within-subject similarity in the functional connectome is not dependent on sex**

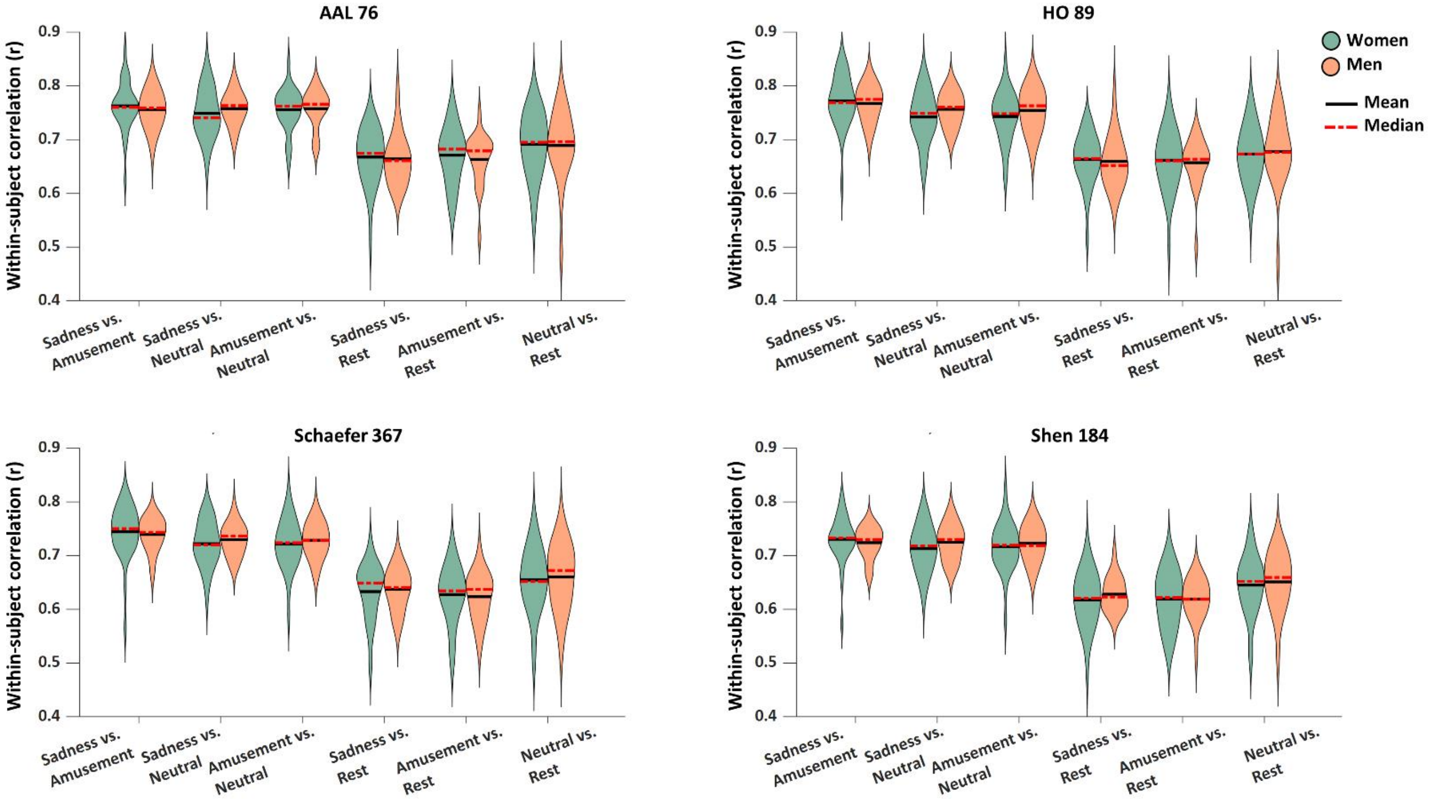


The within-subject similarity in the functional connectome (Pearson's correlation r values) is presented as a function of the pair of brain states. Women are shown in green and men in orange. In each violin, the median is indicated by dashed red lines and the mean by solid black lines. There was no effect of sex or sex-by-state interaction, for any of the parcellation atlases.

**Figure S7. Within-subject similarity in the functional connectome is not dependent on sex: analysis of a balanced subsample of 20 women and 20 men**

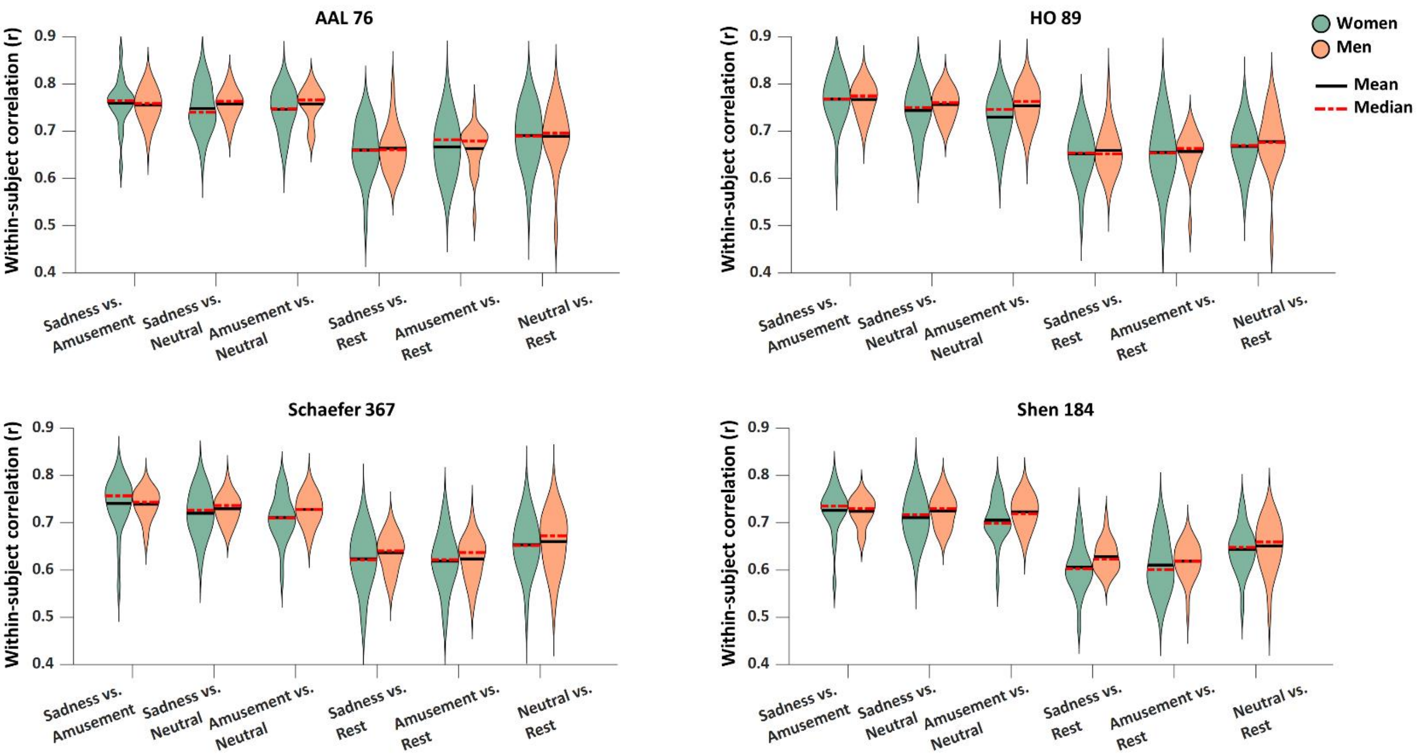


The analysis presented in Figure S6 was repeated with a balanced subsample of 20 women and 20 men. The within-subject similarity in the functional connectome (Pearson's correlation r values) is presented as a function of the pair of brain states. Women are shown in green and men in orange. In each violin, the median is indicated by dashed red lines and the mean by solid black lines. There was no effect of sex or sex-by-state interaction, for any of the parcellation atlases.

**Figure S8. Between-subject similarity in the functional connectome: analysis of a balanced subsample of 20 women and 20 men.**

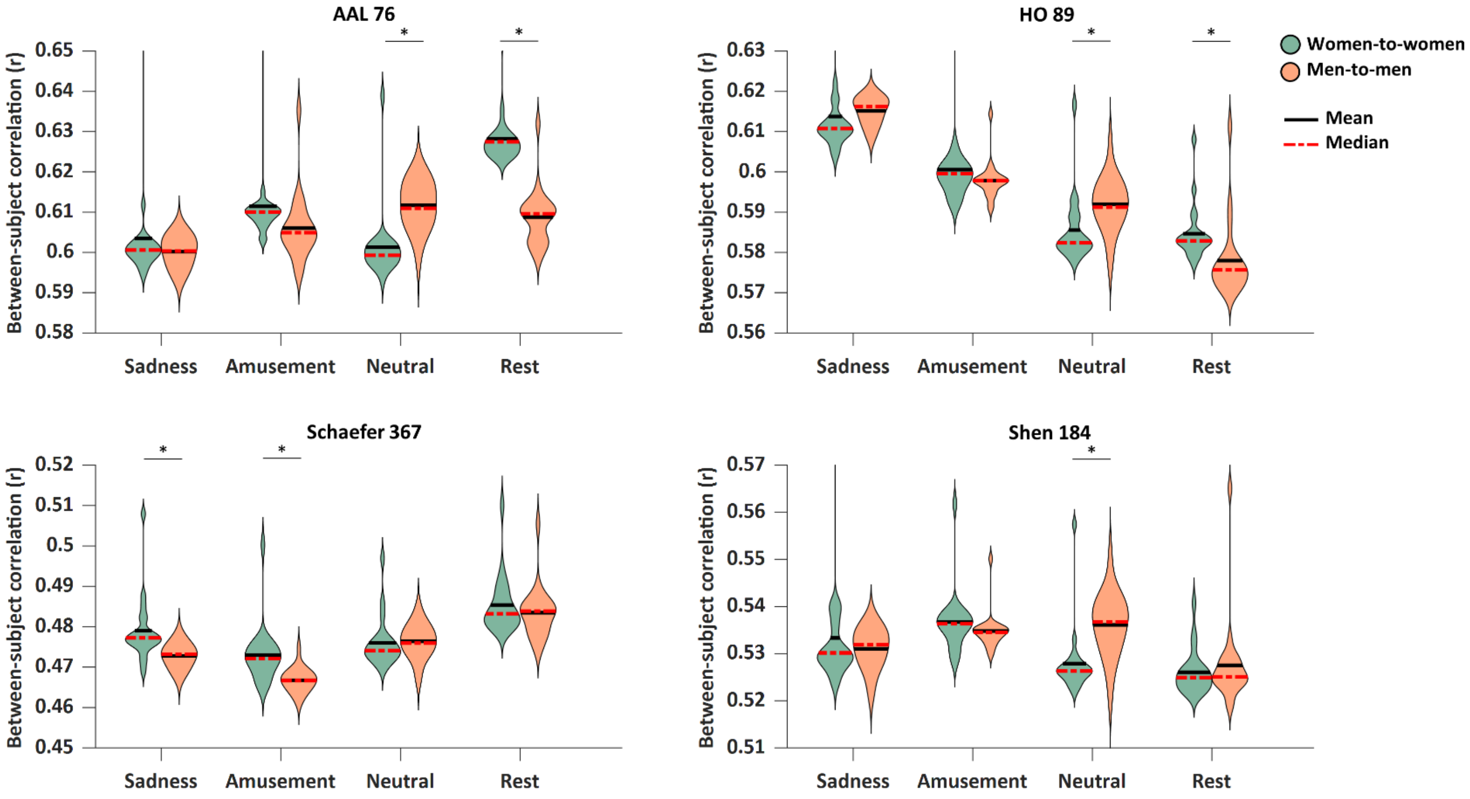


The between-subject similarity in the functional connectome (Pearson's correlation r values) is presented as a function of the brain state, for the four parcellation atlases. The effect of sex was examined by computing women-to-women (green) and men-to-men (orange) similarity. Sex differences varied by state [sex-by-state interaction: AAL: $F(1.82,69.42)=58.57$, $p<10^{-6}$, $\eta_p^2=0.60$; HO: $F(1.76,67.21)=13.15$, $p=3.3\cdot10^{-5}$, $\eta_p^2=0.25$; Shen: $F(1.65,62.75)=12.55$, $p=7.8\cdot10^{-5}$, $\eta_p^2=0.24$; Schaefer: $F(2.38,90.61)=10.02$, $p=4.2\cdot10^{-5}$, $\eta_p^2=0.20$; all Greenhouse-Geisser corrected]. In each violin, the median is indicated by dashed red lines and the mean by solid black lines. Significant differences are marked by asterisks (*), $p<0.05$ Sidak corrected.

**Figure S9. Between-subject similarity in the behavioral report: analysis of a balanced subsample of 20 women and 20 men.**

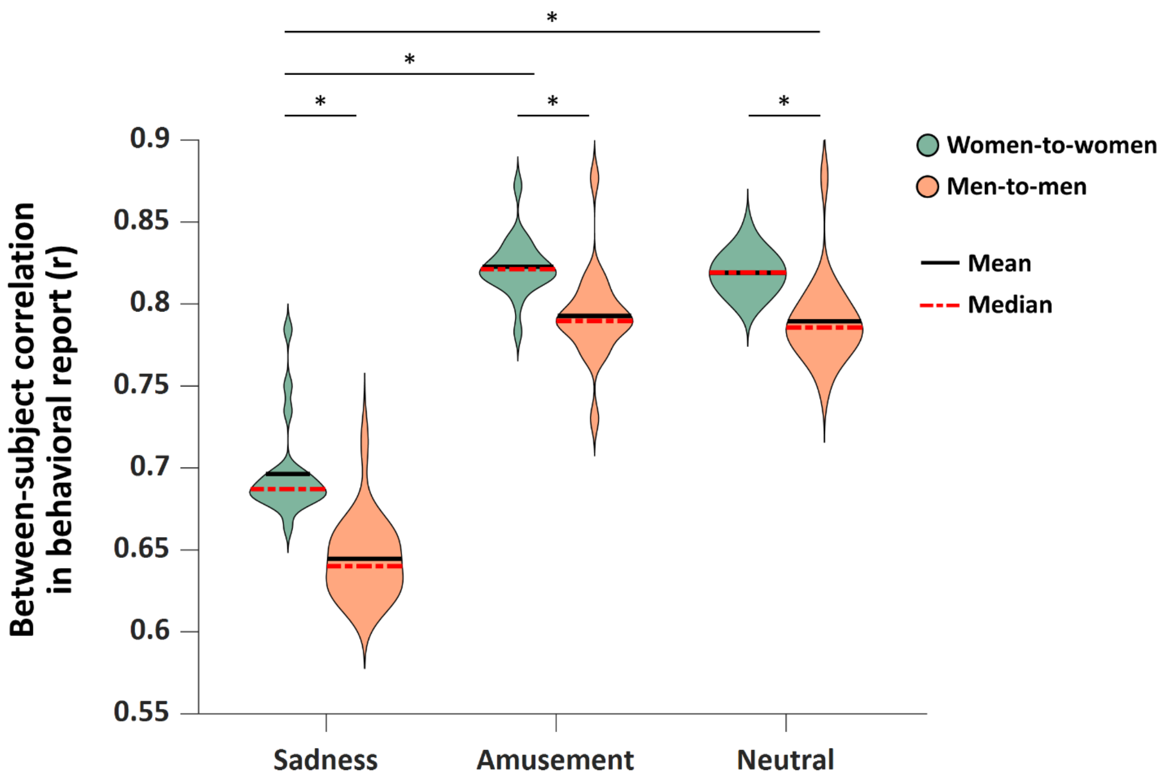


The between-subject similarity in the subjective report of emotional experience is presented as a function of the emotional state. The effect of sex was examined by computing women-to-women (green) and men-to-men (orange) similarity. Women were more similar to other women than men to other men [main effect of sex: $F(1,38)=26.78$, $p<10^{-6}$, $\eta_p^2 = 0.41$. main effect of state: $F(2,76)=794.32$, $p<10^{-6}$, $\eta_p^2 = 0.95$]. There was no sex-by-state interaction. In each violin, the median is indicated by dashed red lines and the mean by solid black lines. Significant differences are marked by asterisks (*), $p<0.05$ Sidak corrected.

**Table S4. The community structure identified for sadness, amusement, neutral, and rest.**

| ROI | Module assignment | | | |
|---|---|---|---|---|
| | Sadness | Amusement | Neutral | Rest |
| Frontal Pole Right | 1 | 1 | 1 | 1 |
| Frontal Pole Left | 1 | 1 | 1 | 1 |
| Insular Cortex Right | 2 | 2 | 2 | 2 |
| Insular Cortex Left | 2 | 2 | 2 | 2 |
| Superior Frontal Gyrus Right | 1 | 1 | 1 | 1 |
| Superior Frontal Gyrus Left | 1 | 1 | 1 | 1 |
| Middle Frontal Gyrus Right | 1 | 1 | 1 | 1 |
| Middle Frontal Gyrus Left | 1 | 1 | 1 | 1 |
| Inferior Frontal Gyrus, pars triangularis Right | 3 | 3 | 3 | 3 |
| Inferior Frontal Gyrus, pars triangularis Left | 3 | 3 | 3 | 3 |
| Inferior Frontal Gyrus, pars opercularis Right | 4 | 4 | 4 | 3 |
| Inferior Frontal Gyrus, pars opercularis Left | 3 | 3 | 3 | 3 |
| Precentral Gyrus Right | 4 | 4 | 2 | 2 |
| Precentral Gyrus Left | 4 | 4 | 2 | 2 |
| Temporal Pole Right | 5 | 3 | 3 | 4 |
| Temporal Pole Left | 5 | 3 | 3 | 4 |
| Superior Temporal Gyrus, anterior division Right | 6 | 3 | 2 | 2 |
| Superior Temporal Gyrus, anterior division Left | 6 | 3 | 2 | 2 |
| Superior Temporal Gyrus, posterior division Right | 6 | 3 | 2 | 2 |
| Superior Temporal Gyrus, posterior division Left | 6 | 3 | 2 | 2 |
| Middle Temporal Gyrus, anterior division Right | 3 | 3 | 3 | 5 |
| Middle Temporal Gyrus, anterior division Left | 3 | 3 | 3 | 5 |
| Middle Temporal Gyrus, posterior division Right | 3 | 3 | 3 | 5 |
| Middle Temporal Gyrus, posterior division Left | 3 | 3 | 3 | 5 |
| Middle Temporal Gyrus, temporooccipital part Right | 3 | 5 | 3 | 6 |
| Middle Temporal Gyrus, temporooccipital part Left | 3 | 3 | 3 | 1 |
| Inferior Temporal Gyrus, anterior division Right | 3 | 3 | 3 | 5 |

| | | | | |
|---|---|---|---|---|
| Inferior Temporal Gyrus, anterior division Left | 3 | 3 | 3 | 5 |
| Inferior Temporal Gyrus, posterior division Right | 1 | 1 | 1 | 1 |
| Inferior Temporal Gyrus, posterior division Left | 1 | 1 | 1 | 1 |
| Inferior Temporal Gyrus, temporooccipital part Right | 4 | 4 | 4 | 6 |
| Inferior Temporal Gyrus, temporooccipital part Left | 4 | 4 | 4 | 6 |
| Postcentral Gyrus Right | 4 | 4 | 4 | 2 |
| Postcentral Gyrus Left | 4 | 4 | 4 | 2 |
| Superior Parietal Lobule Right | 4 | 4 | 4 | 6 |
| Superior Parietal Lobule Left | 4 | 4 | 4 | 6 |
| Supramarginal Gyrus, anterior division Right | 4 | 4 | 4 | 3 |
| Supramarginal Gyrus, anterior division Left | 4 | 4 | 4 | 3 |
| Supramarginal Gyrus, posterior division Right | 1 | 1 | 1 | 3 |
| Supramarginal Gyrus, posterior division Left | 3 | 1 | 1 | 1 |
| Angular Gyrus Right | 1 | 1 | 1 | 1 |
| Angular Gyrus Left | 3 | 3 | 3 | 1 |
| Lateral Occipital Cortex, superior division Right | 7 | 5 | 5 | 6 |
| Lateral Occipital Cortex, superior division Left | 7 | 1 | 5 | 6 |
| Lateral Occipital Cortex, inferior division Right | 7 | 5 | 5 | 7 |
| Lateral Occipital Cortex, inferior division Left | 7 | 5 | 5 | 7 |
| Intracalcarine Cortex Right | 8 | 5 | 6 | 7 |
| Intracalcarine Cortex Left | 8 | 5 | 6 | 7 |
| Frontal Medial Cortex | 1 | 2 | 7 | 4 |
| Juxtapositional Lobule Cortex -formerly Supplementary Motor Cortex- Right | 4 | 4 | 2 | 2 |
| Juxtapositional Lobule Cortex -formerly Supplementary Motor Cortex- Left | 4 | 4 | 2 | 2 |
| Subcallosal Cortex | 1 | 2 | 7 | 4 |
| Paracingulate Gyrus Right | 2 | 2 | 1 | 8 |
| Paracingulate Gyrus Left | 2 | 2 | 1 | 8 |
| Cingulate Gyrus, anterior division | 2 | 2 | 1 | 8 |
| Cingulate Gyrus, posterior division | 1 | 2 | 1 | 5 |
| Precuneous Cortex | 8 | 5 | 1 | 5 |

| | | | | |
|---|---|---|---|---|
| Cuneal Cortex Right | 8 | 5 | 6 | 7 |
| Cuneal Cortex Left | 8 | 5 | 6 | 7 |
| Frontal Orbital Cortex Right | 2 | 2 | 1 | 8 |
| Frontal Orbital Cortex Left | 2 | 2 | 1 | 3 |
| Parahippocampal Gyrus, anterior division Right | 5 | 6 | 7 | 4 |
| Parahippocampal Gyrus, anterior division Left | 5 | 6 | 7 | 4 |
| Parahippocampal Gyrus, posterior division Right | 5 | 6 | 7 | 4 |
| Parahippocampal Gyrus, posterior division Left | 5 | 6 | 7 | 4 |
| Lingual Gyrus Right | 8 | 5 | 6 | 7 |
| Lingual Gyrus Left | 8 | 5 | 6 | 7 |
| Temporal Fusiform Cortex, anterior division Right | 5 | 6 | 7 | 4 |
| Temporal Fusiform Cortex, anterior division Left | 5 | 6 | 7 | 4 |
| Temporal Fusiform Cortex, posterior division Right | 7 | 6 | 5 | 4 |
| Temporal Fusiform Cortex, posterior division Left | 7 | 6 | 5 | 4 |
| Temporal Occipital Fusiform Cortex Right | 7 | 5 | 5 | 7 |
| Temporal Occipital Fusiform Cortex Left | 7 | 5 | 5 | 7 |
| Occipital Fusiform Gyrus Right | 7 | 5 | 5 | 7 |
| Occipital Fusiform Gyrus Left | 7 | 5 | 5 | 7 |
| Frontal Operculum Cortex Right | 2 | 2 | 8 | 3 |
| Frontal Operculum Cortex Left | 2 | 2 | 8 | 3 |
| Central Opercular Cortex Right | 6 | 7 | 2 | 2 |
| Central Opercular Cortex Left | 6 | 7 | 2 | 2 |
| Parietal Operculum Cortex Right | 6 | 7 | 2 | 2 |
| Parietal Operculum Cortex Left | 6 | 7 | 2 | 2 |
| Planum Polare Right | 6 | 7 | 2 | 2 |
| Planum Polare Left | 6 | 7 | 2 | 2 |
| Heschl's Gyrus Right | 6 | 7 | 2 | 2 |
| Heschl's Gyrus Left | 6 | 7 | 2 | 2 |
| Planum Temporale Right | 6 | 7 | 2 | 2 |
| Planum Temporale Left | 6 | 7 | 2 | 2 |
| Supracalcarine Cortex Right | 8 | 5 | 6 | 7 |
| Supracalcarine Cortex Left | 8 | 5 | 6 | 7 |

| | | | | |
|---|---|---|---|---|
| Occipital Pole Right | 7 | 5 | 5 | 7 |
| Occipital Pole Left | 7 | 5 | 5 | 7 |
| Thalamus Right | 9 | 8 | 8 | 8 |
| Thalamus Left | 9 | 8 | 8 | 8 |
| Caudate Right | 9 | 8 | 8 | 8 |
| Caudate Left | 9 | 8 | 8 | 8 |
| Putamen Right | 9 | 8 | 8 | 8 |
| Putamen Left | 9 | 8 | 8 | 8 |
| Pallidum Right | 9 | 8 | 8 | 8 |
| Pallidum Left | 9 | 8 | 8 | 8 |
| Hippocampus Right | 5 | 6 | 7 | 4 |
| Hippocampus Left | 5 | 6 | 7 | 4 |
| Amygdala Right | 5 | 6 | 7 | 4 |
| Amygdala Left | 5 | 6 | 7 | 4 |
| Accumbens Right | 9 | 8 | 8 | 8 |
| Accumbens Left | 9 | 8 | 8 | 8 |
